# *Nepenthes* pitchers *versus* thermogenic flowers: Thermal patterns and their role in prey capture and pollination

**DOI:** 10.1101/2023.09.01.555905

**Authors:** Gokul Baburaj Sujatha, Anil John Johnson, Abdul Azeez Hussain, Sabulal Baby

## Abstract

Prey capture in *Nepenthes* and pollination in angiosperms are two antithetical events; one designed to trap insects and other arthropods (carnivory) and the other to transfer pollen through pollinators (reproduction). In this study infrared thermography is extensively used to obtain thermal profiles of *Nepenthes* pitchers and thermogenic flowers in field conditions. *N. khasiana* pitchers displayed below ambient temperatures during evening-night-morning hours (5 pm to 8-9-10 am); pitcher spots recorded lowest and highest temperatures as 14.4 (6-7 am) and 45.6°C (2 pm), respectively. In the evening-night-morning hours top pitcher spots displayed significant decrease (upto 6.7°C) from the ambient temperature. The average humidities in the night and day periods were 77.15% and 50.15%, respectively. Thermographic tracking of the pitcher (lid) opening in *N. khasiana* demonstrated initial ‘wet’ pitcher in night, which gradually switched to a relatively dry surface in the morning-day times; but the peristome region retained the ‘wetness’ till 10.30 a.m. The prey capturing zones in *Nepenthes* pitchers (peristome, lid and their intersection) are colder favoring prey capture, whereas in thermogenic flowers, floral portions are hotter, providing the thermal requirements for the pollinator. In thermogenic plants, floral zones assist pollination by offering a ‘thermal reward’ through enzymatic processes; but *Nepenthes* traps achieve lower temperature spots by physical (surface microstructures), chemical (extrafloral nectar) and ecological (rain, humidity) factors.

## Introduction

*Nepenthes* pitchers are leaf-evolved biological traps; they entice-trap insects and other arthropods, involve in mutualistic interactions with small mammals and eventually gain nutrients leading to their survival in low supplement growing environments. Carnivory of these pitcher traps is effected through various attraction, deception, capture and digestion strategies (Bohn and Federle, 2004; Kurup et al. 2013; Chen et al. 2016; Baby et al. 2017). *Nepenthes* pitchers consist of distinct zones based on their function, (i) prey capture zone (peristome/lid), (ii) slippery zone (interior waxy region) and (iii) digestive zone (interior bottom portion of digestive fluid) (Bauer and Federle 2009; Wang and Zhou 2016; Baby et al. 2017). The top zone (peristome, lid) of *Nepenthes* pitchers is crucial in attraction and capture of insects and other small organisms. We found UV induced fluorescence emissions primarily from the peristomes of various *Nepenthes* species/hybrids (Kurup et al. 2013). Recent studies have demonstrated fluorescence as a visual signal in floral reproductive structures (Mori et al. 2018), amphibians (frogs), marine organisms, segmented worms, reptiles and other organisms (Taboada et al. 2017; Mori et al. 2018; Kohler et al. 2019; Ben-Zvi et al. 2022). In another recent study, we demonstrated *Nepenthes* pitchers as CO2-encriched cavities, and open pitchers continually emit CO2 attracting preys towards them (Baby et al., 2017). In *Nepenthes*, peristome is a wettable surface with distinct microstructures (Bohn and Federle, 2004; Chen et al. 2016). There are nectaries towards the inner sides of most *Nepenthes* pitcher peristomes. The peristome wettability is through two major factors, extrafloral nectar (EFN) and rain water (Bohn and Federle 2004; Chen et al. 2016); and the wet peristome surface is slippery for the visiting preys. Bauer et al. 2008 reported high rates of prey capture in the evening, night and early morning in *Nepenthes* pitchers. In pollination, a more common natural event, insects are attracted to flowers through various strategies, and among others (pollen, floral nectar (FN)), thermal reward is one of the factors engaging insects to the flowers in several plants (Seymour 1997). Contrastingly, in flowers after visit (pollination) insects ‘fly away’ whereas in *Nepenthes* pitchers the visitors (preys) are destined to ‘fall’ into these traps.

This study is primarily designed to reflect on the influence of thermal patterns in prey capture (pitchers) in *Nepenthes* and pollination (flowers) in angiosperms. Infrared radiation is emitted by any object at temperatures above absolute zero, and infrared thermography is a technique which detects infrared energy emitted from the object, converts it to temperature, and displays the image of temperature distribution. It is a non-invasive technique and requires no contact to measure the temperature accurately (Merlot et al. 2002; Wang et al. 2004; Dyer et al. 2006; Usamentiaga et al. 2014; Tattersall et al. 2016; Harrap et al. 2017). Here we report (i) the thermal profiles of *Nepenthes* pitchers in field and lab conditions, (ii) their comparison with thermal patterns of thermogenic flowers and (iii) discuss the influence of thermal patterns in prey capture (carnivory) and pollination (reproduction). This is the first study reporting thermal profiles of pitcher regions of various *Nepenthes* species/hybrids and correlating these patterns to relative humidity and atmospheric temperature. Moreover, the study records the periodical changes in thermal patterns in *Nepenthes* pitcher (lid) opening in field conditions.

## Materials and methods

### Study species

Pitchers of *Nepenthes khasiana* Hook. f. (Fig. 1) and *Nepenthes* hybrids (Fig. S1) and flowers of *Victoria amazonica* (Poepp.) Klotzsch (family: Nymphaeaceae, common name: giant water lily or Amazon water lily) (Fig. S4) and *Nelumbo nucifera* Gaertn. (Nelumbonaceae, sacred or Indian lotus) (Fig. S6) were sampled from the garden sites of Jawaharlal Nehru Tropical Botanic Garden and Research Institute (JNTBGRI), Palode, Kerala in south India.

**Fig. 1.**
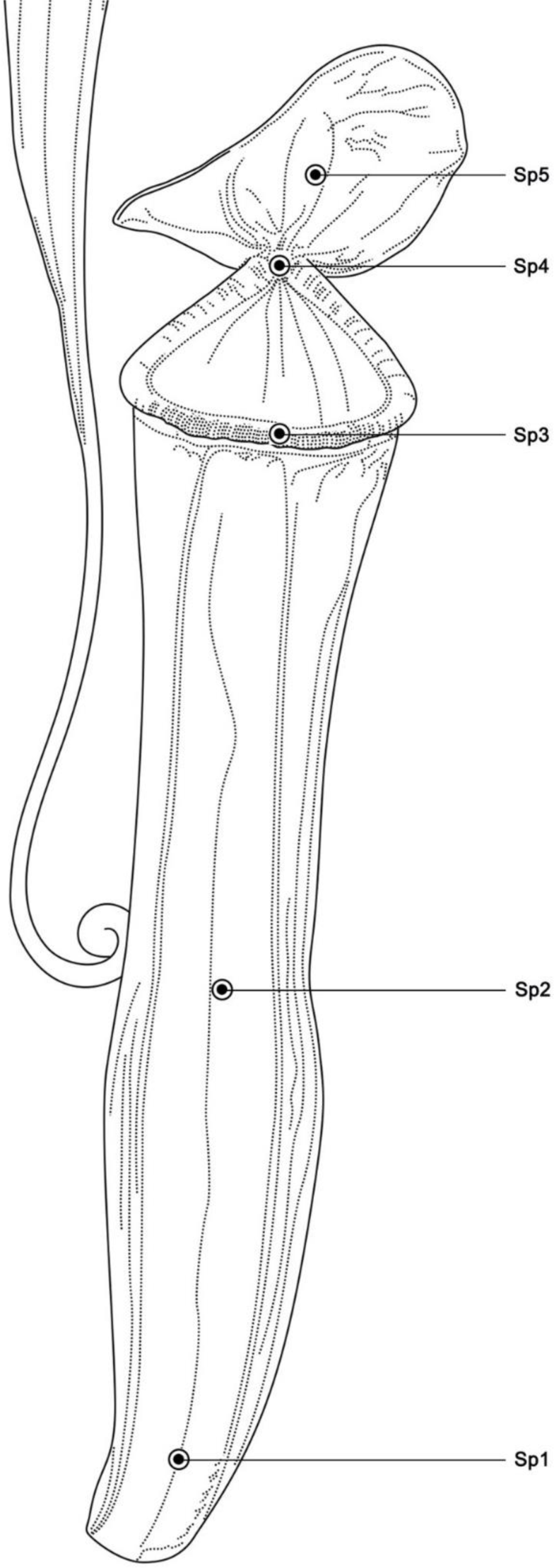
Line diagram of *N. khasiana* open pitcher with spots sp1 to sp5 marked from bottom to top. In unopen pitchers, only three spots (sp1 to sp3) were marked (measured), with sp3 as the front centre spot of the top peristome-lid interface (in closed position).

### Infrared thermography

*Nepenthes khasiana*, *Nepenthes* hybrid pitchers and thermogenic flowers were subjected to infrared thermography in the field and their thermal profiles were recorded using infrared camera (FLIR 420bx, USA) with FOL lens (18 mm), IR pixel resolution (240 x 320), temperature range -20 to 350°C and thermal sensitivity < 0.045°C. The thermal patterns were analyzed using FLIR Tools Version 5.5. Infrared thermographic profiles of pitchers and flowers were recorded in the field in 1 hour intervals for 24 or 48 hours (day & night) (Supporting Information). *N. khasiana* pitchers were also thermographed in the lab under controlled conditions (temperature: 22°C, humidity: 60%). Temperatures of 3 spots (sp1, sp2, sp3) in *N. khasiana* unopen pitchers (P1U-P6U), 5 spots (sp1, sp2, sp3, sp4, sp5) in open pitchers (P1O-P6O) and chosen spots on thermogenic flowers were recorded from their thermal images (Figs. 1-3, S2-S7). Infrared thermographic measurements of unopen/open *N. khasiana* pitchers (P1U-P3U & P1O-P3O) were recorded in January 2019 and (P4U-P6U & P4O-P6O) in March 2021 (Supporting Information). Plant specimens not under direct sunlight were chosen for these measurements. All thermographic measurements in the field were carried out on non-rainy conditions. Leaf specimens of attached to *N. khasiana* and *Nepenthes* hybrid pitchers were also recorded, but no specific patterns of interest were observed. An Environmental Meter (Extech Instruments 45170, FLIR Systems, India) was used to measure the relative humidity and atmospheric temperature.

**Fig. 2.**
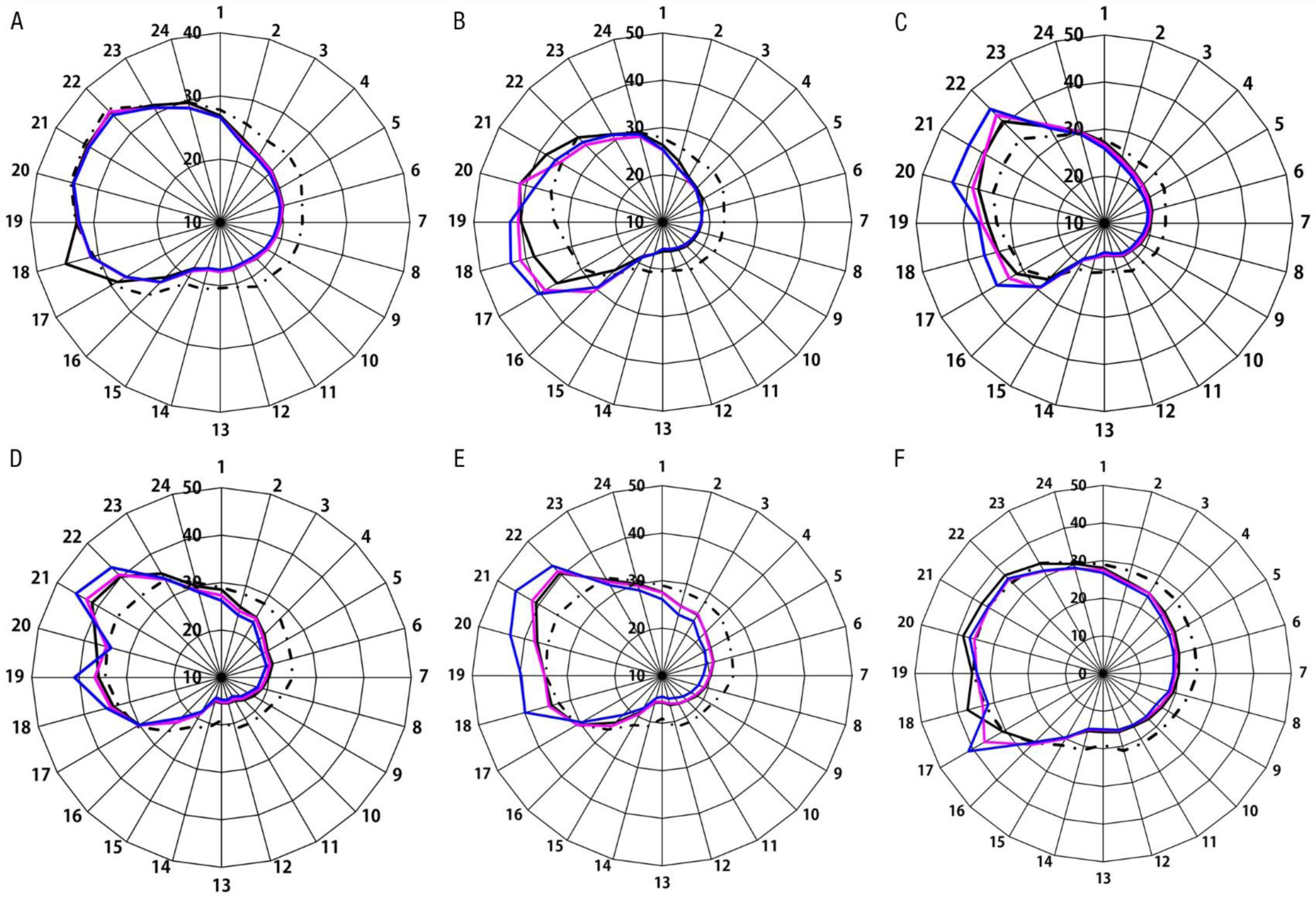
Infrared thermographic profiles of six unopen *N. khasiana* pitchers in field conditions; (A) P1U, (B) P2U, (C) P3U, (D) P4U, (E) P5U, (F) P6U. P1U to P3U & P4U to P6U were recorded in January 2019 and March 2021, respectively (dotted line: ambient atm. temp., black: sp1, pink: sp2; blue: sp3; spots are as labelled in Fig. 1; X-axis (circular) (A-F): 1-24 are 24 h time at 1 h measurement intervals starting 6 pm to 5 pm, next day (Tables S1-S6); Y-axis: temperature in degrees).

**Fig. 3.**
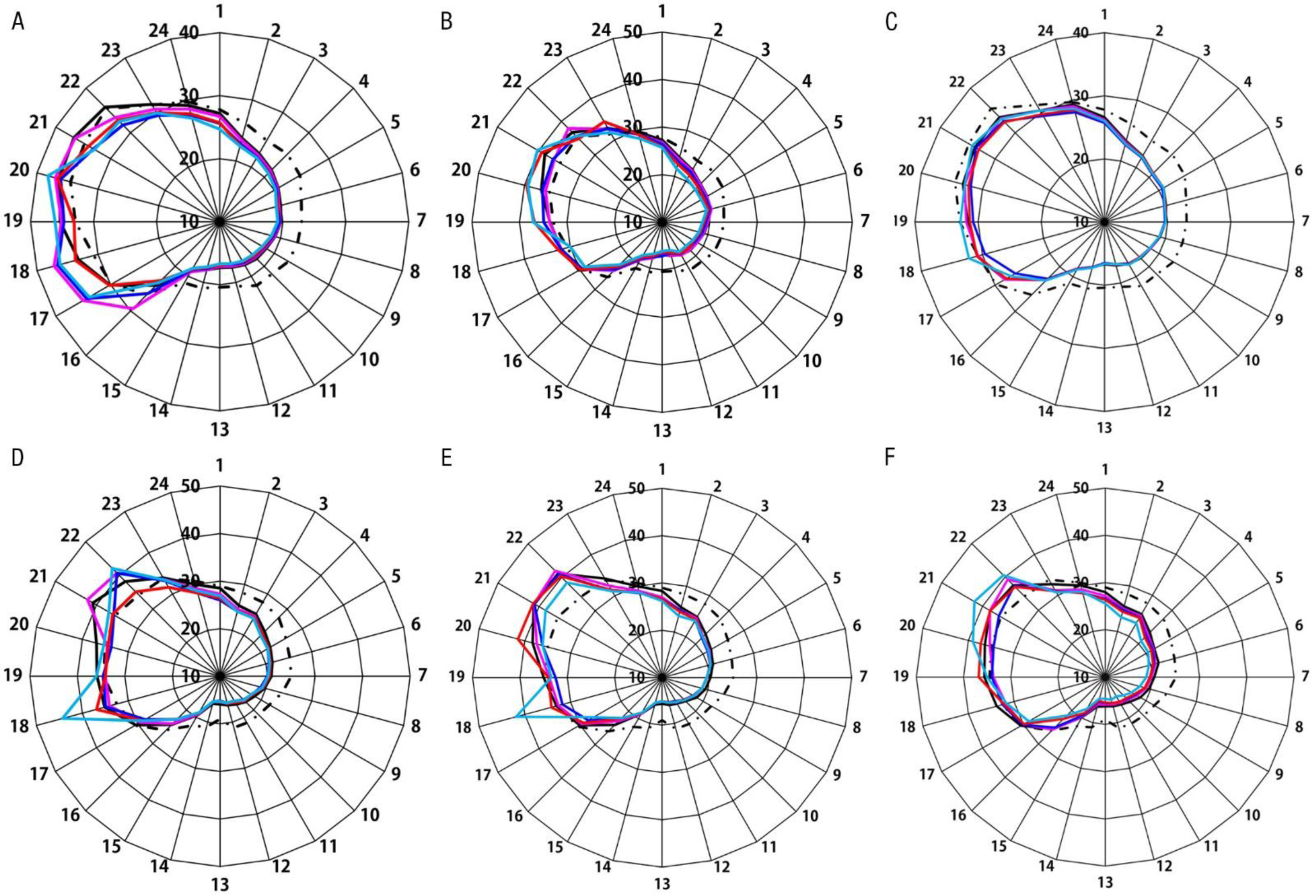
Infrared thermographic profiles of six open *N. khasiana* pitchers in field conditions; (A) P1O, (B) P2O, (C) P3O, (D) P4O, (E) P5O, (F) P6O. P1O to P3O & P4O to P6O were recorded in January 2019 and March 2021, respectively (dotted line: ambient atm. temp., black: sp1, pink: sp2; blue: sp3, red: sp4, green: sp5; spots are as labelled in Fig. 1; X-axis (circular) (A-F): 1-24 are 24 h time at 1 h measurement intervals starting 6 pm to 5 pm, next day (Tables S7-S12); Y-axis: temperature in degrees).

### Scanning electron microscopy

Scanning electron microscopy (SEM) imaging of *N. khasiana* sections was carried out on a SEM (Hitachi S-2400, Japan). *N. khasiana* pitcher sections were fixed in 3% glutaraldehyde in phosphate buffer and kept overnight. Sections were then dehydrated sequentially with different concentrations of ethanol (30%, 50%, 70% 15 min each, two changes and 90% and 100% ethanol, 35 min each, two changes). The dehydrated sections were then subjected to critical point drying, gold coated and viewed on the SEM. Again, *N. khasiana* peristome and lid samples were mounted over aluminium stub using double sided carbon tape and imaged using high resolution direct SEM (FEI Quanta 200, Netherlands).

## Results

### Nepenthes pitchers, thermal patterns

Thermal profiles of three spots (sp1, sp2, sp3; spots labelled in Fig. 1) in unopen *N. khasiana* pitchers (P1U to P6U) (Figs. 1, 2) and five spots (sp4: peristome-lid joint and sp5: centre of the open lid; spots labelled in Fig. 1) in open pitchers (P1O to P6O) (Figs. 1, 3) were measured in field and laboratory conditions. The two top spots (sp4, sp5) are exposed (or measurable) only in open pitchers. *N. khasiana* pitchers showed temperature zones at its various regions (spots). For example, in the unopened pitcher (P2U) (Fig. 2B) from 5 pm to 8 am (average humidity and temperature: 79.4 ± 4.42% and 23.54 ± 2.68°C), pitcher temperature zones (sp1-sp3) remained below the atm. temperature and from 9 am to 4 pm (average humidity and temperature: 56.01 ± 13.13% and 31.98 ± 3.07°C) pitcher zones exceeded the ambient temperature. Among the pitcher zones, sp3 (blue, pitcher top) attained a maximum temperature of 43.3°C at 11 am, and sp2 (violet) and sp3 (blue) attained the lowest temperatures of 15.7°C at 6 am. Maximum temperature difference between the ambient temperature and pitcher zones (sp1, sp2, sp3) in the nighttime (5 pm to 8 am) was (24.7-18.9°C (sp3)) 5.8°C at 10 pm. Similar temperature difference in daytime (9 am to 4 pm) was (43.3 (sp3) - 30.7°C) a significant 12.6°C (above ambient). In the nighttime, all three pitcher spots (sp1, sp2, sp3) are almost overlapping throughout, but in daytime spots (pitcher regions) are distinct, with maximum temp difference of 43.3 (sp3) - 38.2°C (sp1), i.e., 5.1 degrees at 11 am. The top spot (sp3) of *Nepenthes* unopened pitchers was the hottest.

In an example of an open *N. khasiana* pitcher (P1O) (Fig. 3A), the nighttime zone extended from 5 pm to 8 am (morning) next day. In this time range, the 5 pitcher spots (sp1-sp5) showed temperatures below the atmospheric temperatures. Ambient temp. range (5 pm to 8 am) was from 20.4-29.7°C with an average 23.54 ± 2.68°C and humidity ranged from 67.5-83.5% (average, 79.40 ± 4.42%). Lowest pitcher temp. (spot) in the night zone was 16.7°C (6 am, sp5) and highest temp. was 29.1°C (5 pm, sp1). In the daytime zone (9 am to 4 pm), most pitcher spots (zones) exceeded the ambient temperature. Ambient temp. and humidity ranges were 26.3-35.4°C (average, 31.98 ± 3.07°C) and 41.5-79.0% (56.01 ± 13.13%), respectively. Lowest pitcher temp (spot) in the day zone was 23.2°C (9 am, sp4) and highest was 38.2°C (1 pm, sp5).

In open (P1O to P6O) and unopen (P1U to P6U) pitchers (6 each) monitored for 48 h in the field (Figs. 2, 3), the pitcher temperatures were lower than the atmospheric temperatures starting evening (5 pm), throughout the night (6 pm to 6 am), and in the morning hours (8 am or 9 am or 10 am). These lower temperatures of the pitcher zones extended to daytime (up to 10 am), depending on the type of the pitchers (open/unopen) and their ecological conditions. In six each unopen and open pitchers, spots (sp1-sp3 & sp1-sp5) remained > 3 degrees below the atmospheric temperatures in the evening-night-morning hours (Tables S1-S12, Figs. S2, S3); and the pitcher spots (unopen pitchers: sp1-sp3; open pitchers: sp1-sp5) were mostly overlapping. The atmospheric-spot temperature difference in the night time was 6.7°C (each) in the unopen pitcher P4U (sp3, 4 am) and open pitcher P6O (sp5, 9 pm).

The lowest temperatures in six unopened (P1U-P6U) pitchers were 17.3 (P1U: 4 am, sp3), 15.7 (P2U: 6 am, sp2/sp3), 16.2 (P3U: 6 am, sp3), 14.4 (P4U: 7 am, sp3), 14.4 (P5U: 6 am, sp3), 14.8 (P6U: 6 am, sp3) degrees and in six opened (P1O-P6O) pitchers were 16.7 (P1O: 6 am, sp5), 16.1 (P2O: 5-6 am, sp5), 16.5 (P3O: 6 am, sp3), 15.2 (P4O: 6 am, sp3), 15.0 (P5O: sp2-sp5, 6 am), 14.6 (P6O: 6 am, sp5), respectively. So, in the measurement conditions, *N. khasiana* pitchers recorded lowest temperatures in the range 14.4-17.3 degrees at 4-7 am. In all six unopened pitchers (P1U-P6U) sp3 showed the lowest temperatures (14.4-17.3 degrees) in the morning hours. In opened pitchers (P1O-P6O), sp3 & sp5 recorded the lowest temperatures (14.6-16.7 degrees) under the measurement conditions.

Similarly, the highest temperatures in unopened pitchers (P1U-P6U) were 35.3 (P1U: 11 am, sp1), 43.3 (P2U: 11 am, sp3), 44.3 (P3U: 3 pm, sp3), 45.4 (P4U: 2 pm, sp3), 45.6 (P5U: 2 pm, sp3), 41.3 (P6U: 10 am, sp3). Open pitchers (P1O-P6O) recorded highest temperatures as: 38.2 (P1O: 1 pm, sp5), 40.3 (P2O: 2 pm, sp5), 34.2 (P3O: 2 pm, sp5), 44.3 (P4O: 11 am, sp5), 41.9 (P5O: 11 am, sp5 & 3 pm, sp2), 42.0 (P6O: 2 pm, sp5). The highest pitcher temperatures (34.2-45.6 degrees) were registered from 10 am-3 pm. In 6 unopened pitchers (P1U-P6U), sp3 (5) & sp1 (1) recorded highest temperatures (35.3- 45.6 degrees). In 6 open pitchers (P1O-P6O), sp5 (6) & sp2 (1) recorded highest temperatures (34.2-44.3 degrees).

In unopen/open pitchers (P1O-P6O & P1U-P6U) in 24 h (under the field experimental conditions), the lowest and highest temperatures between pitcher spots varied markedly, unopen pitchers: (P1U: 35.3 (sp1)-17.3 (sp3) = 18°C; P2U: 43.3 (sp3)-15.7(sp2/sp3) = 27.6°C; P3U: 44.3 (sp3) - 16.2 (sp3) = 28.1°C; P4U: 45.4 (sp3)-14.4 (sp3) = 31°C; P5U: 45.6 (sp3)-14.4 (sp3) = 31.2°C; P6U: 41.3 (sp3)-14.8(sp3) = 26.5°C); open pitchers: (P1O: 38.2 (sp5)-16.7 (sp5) = 21.5°C; P2O: 40.3 (sp5)-16.1 (sp5) = 24.2°C; P3O: 34.2 (sp5)-16.5 (sp3) = 17.7°C; P4O: 44.3 (sp5)-15.2 (sp3) = 29.1°C; P5O: 41.9 (sp5)-15.0 (sp2-sp5) = 26.9°C; P6O: 42.0 (sp5)-14.6 (sp5) = 27.4°C). Unopen and open pitchers (6 each) in 24 hours in field measurements showed maximum variations of 31.2 and 29.1°C, respectively, between their spots.

In unopen pitchers (P1U-P6U), the temperature variations (lowest/highest) of pitcher spots against atmospheric temperature (of the respective spot/time): (P1U: low, 4.4°C, 10 pm, 2 am, 4 am (sp3), high, 4.6°C, 11 am (sp1); P2U: low, 5.8°C, 10 pm (sp3), high, 12.6°C, 11 am (sp3); P3U: low, 4.8°C, 10 pm (sp3), high, 8.9°C, 1 pm, 3 pm (sp3); P4U: low, 6.7°C, 4 am (sp3), high, 9.9°C, 2 pm (sp3); P5U: low, 6.5°C, 9 pm (sp3), high, 10.1°C, 2 pm (sp3); P6U: low, 6.3°C, 12 am (sp3), high, 10.6°C, 10 am (sp3)). Similar variations in open pitchers (P1O-P6O): (P1O: low, 4.8°C, 2 am (sp5), high, 6.5°C, 11 am (sp2); P2O: low, 5.4°C, 9 pm, (sp5); high, 5.5°C, 4 pm (sp5); P3O: low, 4.4°C, 9 pm (sp5), high, 3.0°C, 3 pm (sp4); P4O: low, 5.5°C, 7 am (sp2, sp3, sp4, sp5), high 11.8°C, 11 am (sp5); P5O: low, 5.8°C, 5 am (sp4), high, 9.4°C, 11 am (sp5); P6O: low, 6.7°C, 9 pm (sp5), high, 6.5°C, 2 pm (sp5). *Nepenthes* hybrid pitchers (open and unopen) also showed similar thermographic patterns (Fig. S1).

In January 2019 measurements (P1U to P3U & P1O to P3O), average humidity and ambient temperature from 5 pm to 9 am were 79.38% and 23.71 degrees, respectively (Tables S1-S3, S7-S9, Figs. S2, S3). Average humidity and ambient temperature from 10 am to 4 pm were 52.73% and 32.79 degrees. Similarly, in March 2021 (P4U to P6U & P4O to P6O), average humidity and ambient temperature from 5 pm to 9 am were 74.91% and 24.54 degrees whereas these parameters from 10 am to 4 pm were 47.56% and 33.74 degrees, respectively (Tables S4-S6, S10-S12, Figs. S2, S3).

In our infrared thermographic observations, we monitored the opening of one *N. khasiana* pitcher (Fig. 4) in the field. The initiation of pitcher (lid) opening was in the early morning hours (4.30 am) (Fig. 4B), and at this point the pitcher and linked tendril/lamina were wet throughout, displaying blue colour in infrared thermography (with maximum colour intensity at the peristome/lid interface). The lid opening extended to the day zone (10.30 am), and the blue (wet) ring was retained at the peristome-lid interphase till 10.30 am (Fig. 4H). Most of the pitcher surface (and the tendril/lamina linked to the pitcher) became hotter (gradually turned yellow) by 8.30 am (Fig. 4F), and at 11.30 am the entire pitcher became almost uniformly hot (yellow) (Fig. 4I). This pattern clearly indicates the moisture-retaining capacity of the peristome-lid interphase (Fig. 4F-H) in *Nepenthes* pitchers.

**Fig. 4.**
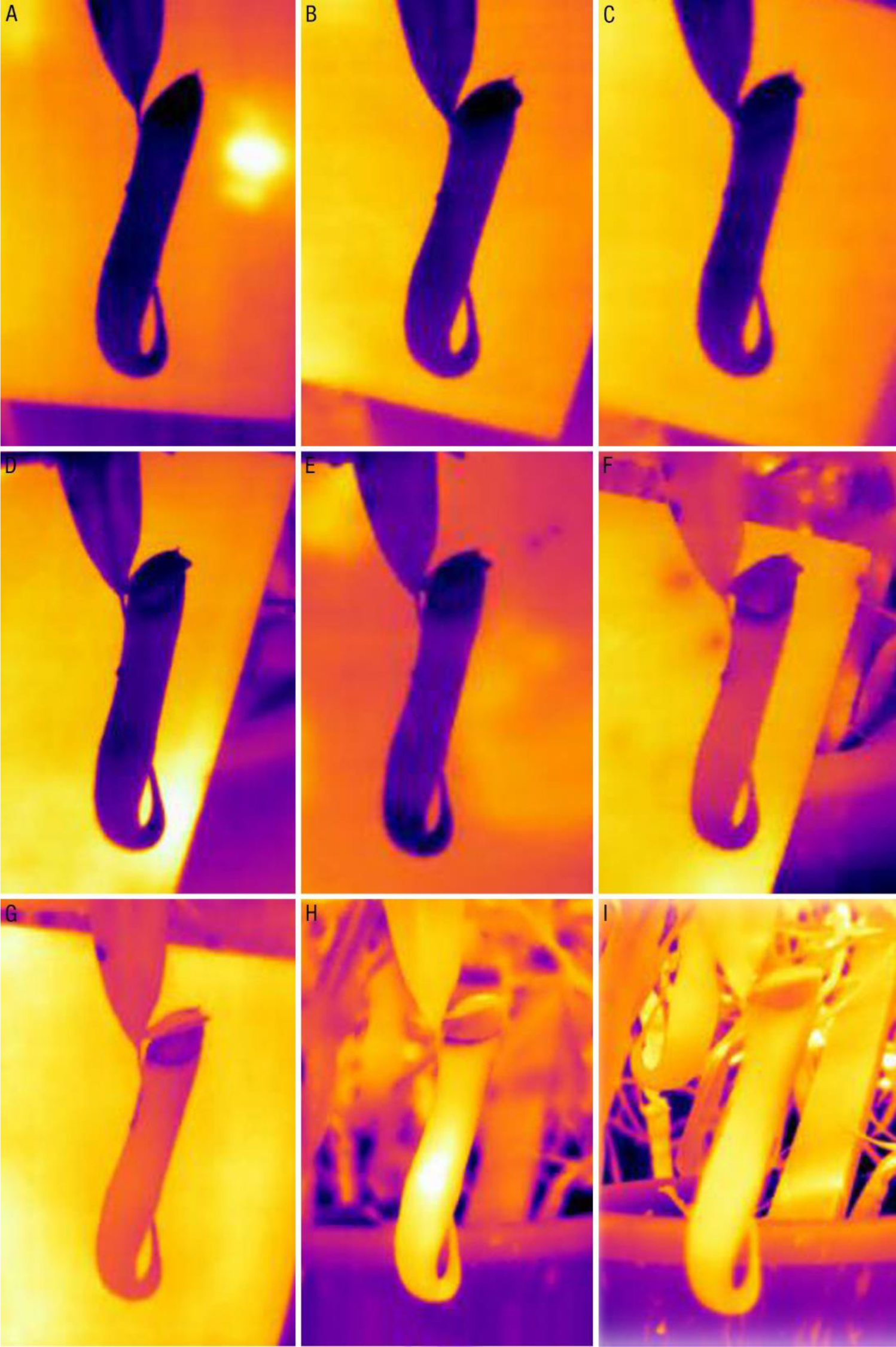
*N. khasiana* pitcher opening monitored by infrared thermography, (A) 12.30 am, (B) 4.30 am, (C) 5.30 am, (D) 6.30 am, (E) 7.30 am, (F) 8.30 am, (G) 9.30 am, (H) 10.30 am, (I) 11.30 am, measurements on March 28, 2018.

The variation in pitcher temperatures over time (of the day) and between the pitcher spots is largely influenced by the ecological parameters (ambient temp., humidity, wind, rain etc.). To clarify this, we measured the thermal patterns of *N. khasiana* pitchers in a controlled atmosphere, at stable temperature of (22°C) and humidity (60%). A *N. khasiana* potted plant with multiple pitchers was brought to a laboratory room with controlled atmosphere (22°C/60%), allowed to stabilize within the room and then the temperature patterns at five spots (sp1-sp5) in a mature open pitcher were measured. In these conditions, all five spots (sp1 to sp5) exceeded the ambient temp. (22°C), with the lowest temp. 22.4°C (sp4) and highest temp. 24.5°C (sp2 & sp5) (Fig. 5). sp3 (average 23.06°C) and sp4 (22.74°C) stayed as the lowest temperature spots throughout the measurements under these controlled conditions. The differences between spots (average temp., sp1 to sp5) and ambient (22°C) temperatures were 2.08°C (sp1), 2.18°C (sp2), 1.06°C (sp3), 0.75°C (sp4) and 2.16°C (sp5) (Fig. 5). On spraying water uniformly, the entire pitcher turned blue (indicating low temperatures); air flow also had a role on the surface temperatures of pitchers.

**Fig. 5.**
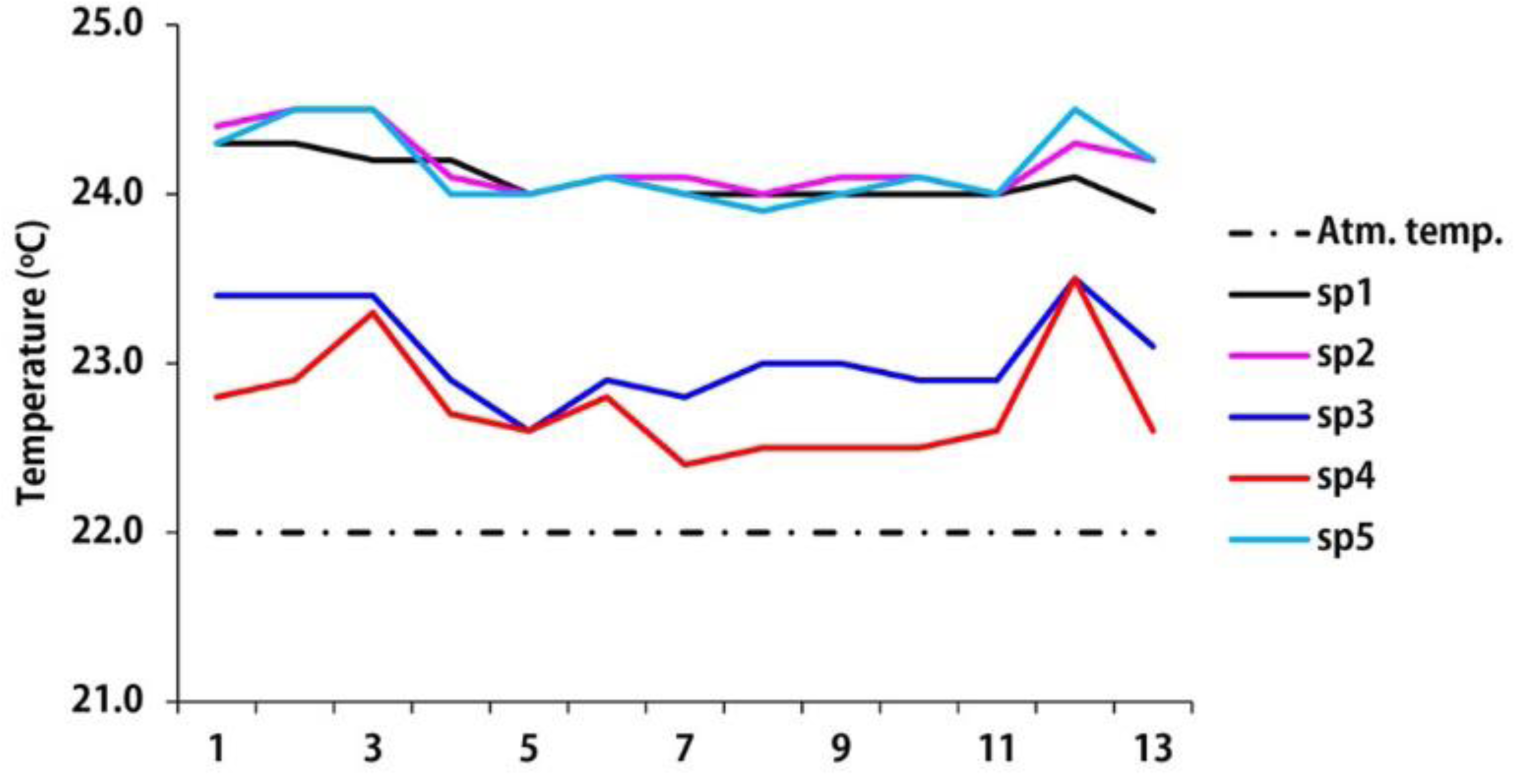
Infrared thermographic profile of *N. khasiana* pitcher in controlled laboratory conditions; (dotted line: ambient atm. temp., black: sp1, pink: sp2; blue: sp3, red: sp4, green: sp5; spots are as labelled in Fig. 1; X-axis: measurement data points (13) in 30 min duration).

### Thermogenic flowers, thermal patterns

The flowers of two known thermogenic plants, *V. amazonica* and *N. nucifera*, in the same field were subjected to infrared thermographic analysis. In *V. amazonica* flower (second day of flowering, 4 pm to 10 am, 18 h, 19 readings, January 2019), the floral (outer) base portion (sp2) is warmer than atm. temp., and the central portion of the flower (upper floral lobes, sp3) and the base lobes (sp1) remained mostly below atm. temperatures (Figs. 6A, S5, Table S13). In a second set of measurements (two days of flowering, 4 pm to 3 pm, 48 h, January 2022), *V. amazonica* flower bud was fully visible (above the water level) at 16.00 pm in the evening. At 17.00 pm, the hot spot (ring) appeared and the flower (white) completely bloomed by 19.00 pm; and at this stage, all three zones (sp1, sp2, sp3) were clearly visible. After midnight, the temperature of the hottest floral spot (sp2) was lower than the atmospheric temperature, and at early morning sp2 temperature increased. After 11 am till afternoon the atmospheric temperature was higher than the floral spots and the flower turned pink in colour. Along with colour change in the morning hours the flower began to close partially and opened again in the afternoon. This night and day temperature changes continued in the second day. The outer floral petals began to partially drown in the water, the hot ring faded in the second day night and the central portion of the flower began to appear. In the second day morning, the flower was seen floating over the water, differentiation of the spots was difficult and the observations ended at 15.00 pm. In these measurements (two days of flowering), sp2 was mostly hotter, whereas sp1 and sp3 stayed below the ambient atmospheric temperatures (Figs. 6B, S5, Table S14).

**Fig. 6.**
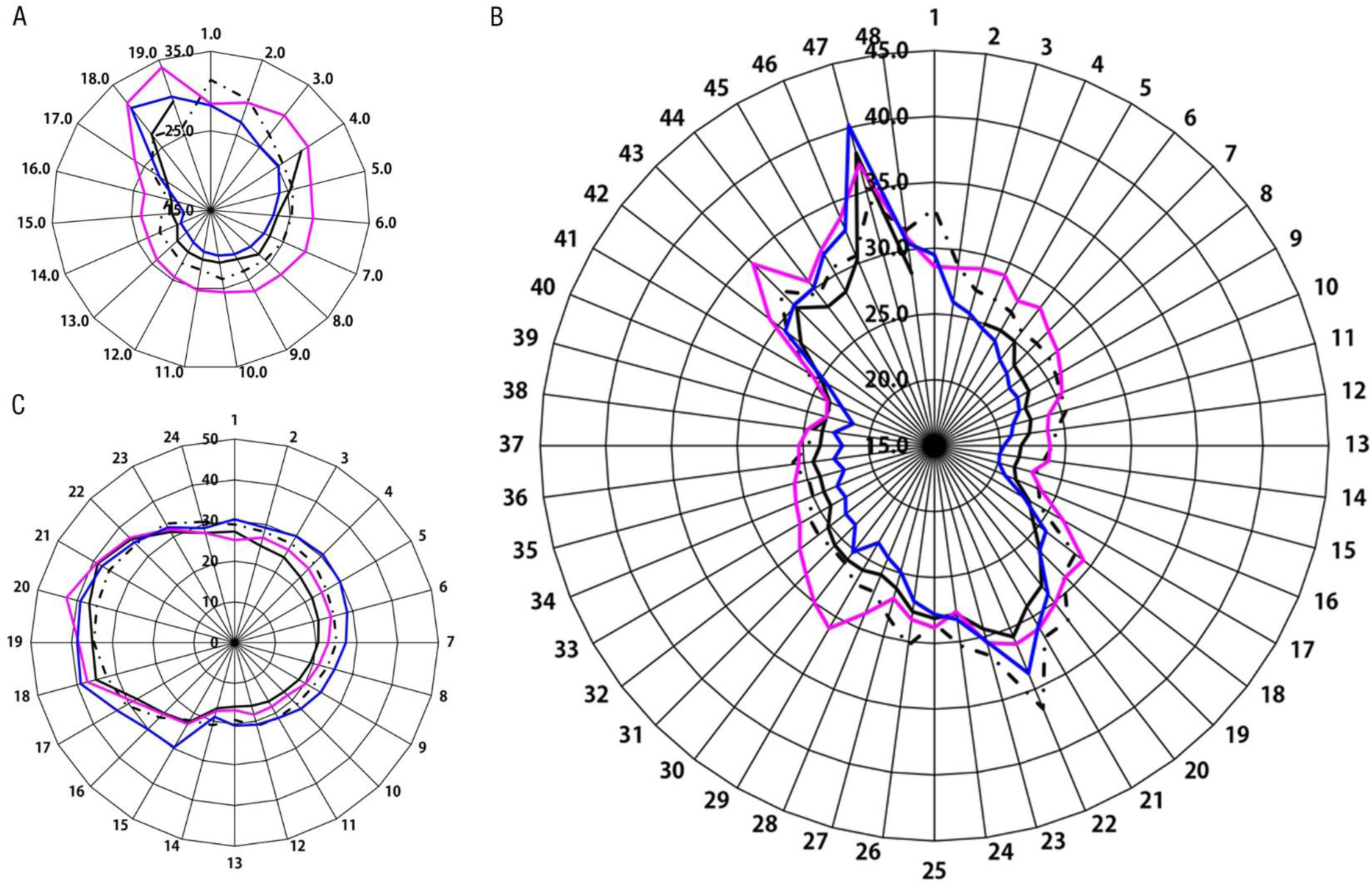
Thermographic patterns of (A) *V. amazonica*, 4 pm to 10 am (18 h), January 2019; (B) *V. amazonica*, 4 pm to 3 pm (48 h), March 2021; (C) *N. nucifera*, 6 pm to 5 pm (24 h), March 2021; (dotted line: ambient atm. temp., black: sp1, pink: sp2; blue: sp3; X-axis (circular) (A): 1-19, time at 1 h measurement intervals starting 4 pm to 10 am, next day (Table S13); X-axis (circular) (B): 1-48, time at 1 h measurement intervals starting 4 pm to 3 pm, 48 h (Table S14); X-axis (circular) (C): 1-24, time at 1 h measurement intervals starting 6 pm to 5 pm, 24 h (Table S15); Y-axes (A-C): temperature in degrees).

In *N. nucifera* (24 h, March 2021) the interior spot (sp3) was warmer compared to the two outer spots (sp2 and sp1); sp3 is higher than ambient temperatures throughout the 24 h field measurements (Figs. 6C, S6, Table S15). sp1 and sp2 remained below ambient temperatures from 4 pm to 10 am. Even in these time periods (evening-night-morning), sp3 retained higher temperatures compared to sp1, sp2 and ambient temperatures. In the 24 h measurements, average sp1, sp2 and sp3 temperatures were 24.93°C, 26.55°C and 29.36°C (average humidity 66.93%, ave. atmospheric temp. 27.22°C). The lowest floral temp. was 15.7°C (sp1, 6.30 am) and at this same time sp2 and sp3 recorded 16.6 and 20.4°C (sp3-sp1, 4.7°C); the highest floral temp. was 42.7°C (sp2, 1.30 pm) and sp1 and sp3 at 1.30 pm were at 36.9 and 39.2°C, respectively (sp2-sp1, 5.8°C).

### SEM of *N. khasiana* pitcher

SEM images displayed glands at the inner sides of pitcher lids, nectaries at the inner sides of peristome, modified stomata at the waxy zone (below peristome) and secretory glands at the digestive zone of *N. khasiana* pitchers (Fig. 7). Direct SEM displayed nectar deposition and glands on pitcher lids (Fig. 7).

**Fig. 7.**
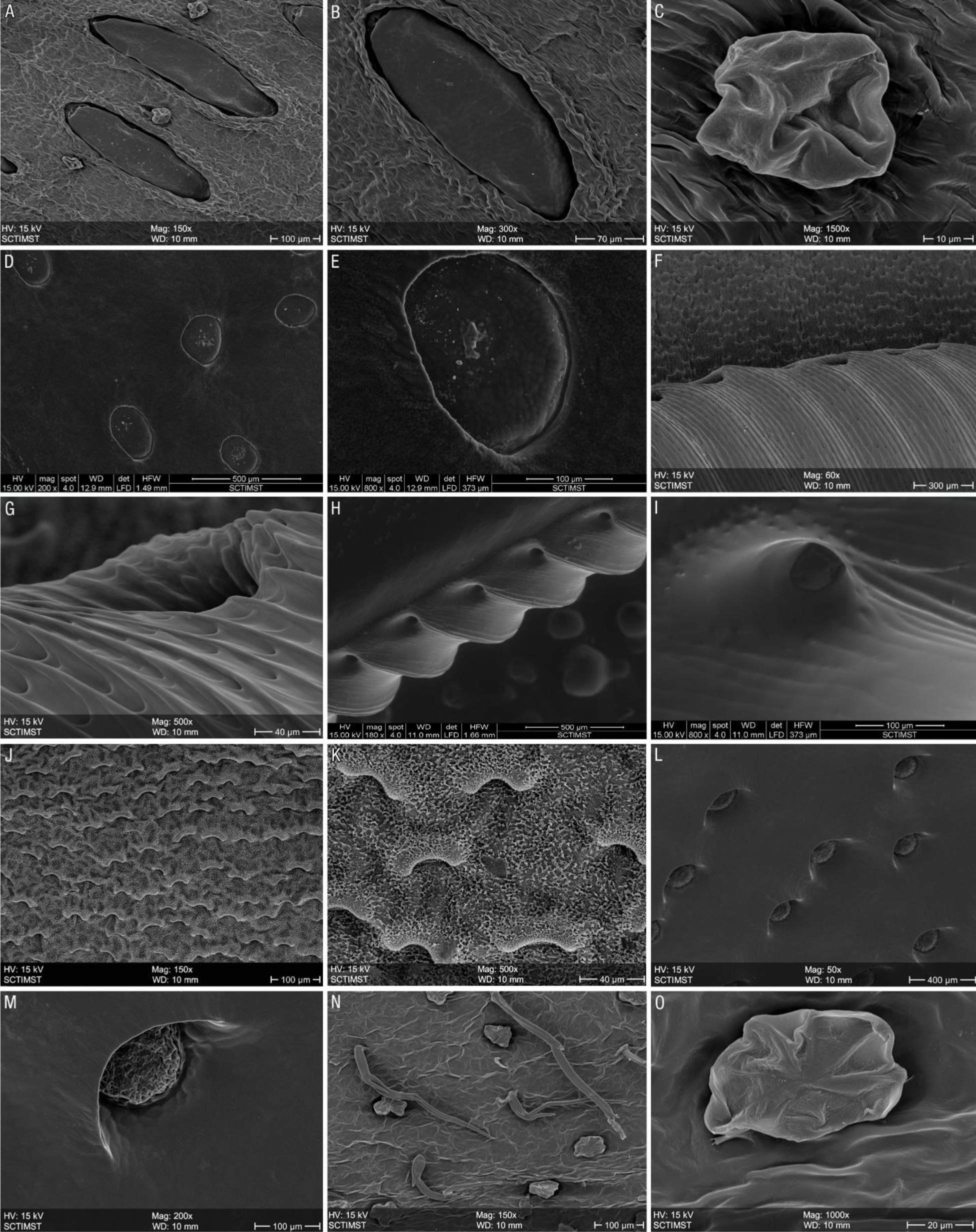
SEM pictures of various regions of *N. khasiana* pitchers, (A) lid, inner side; (B) lid, inner side, gland (expansion); (C) lid, inner side, gland (expansion); (D) direct SEM, lid, inner side, nectar deposition; (E) direct SEM, lid, inner side, nectar deposition (expansion); (F) peristome, nectaries at the inner side; (G) peristome, nectary (expansion); (H) direct SEM, peristome, nectaries at the inner side; (I) direct SEM, peristome, nectary (expansion); (J) slippery (waxy) zone; (K) slippery (waxy) zone (expansion); (L) liquid zone, glands; (M) liquid zone, glands (expansion); (N) pitcher, outer side, glands; (O) pitcher, outer side, gland (expansion).

## Discussion

### Nepenthes pitchers, thermal patterns

In *N. khasiana* open (P1O to P6O) and unopen (P1U to P6U) field pitchers (Figs. 2, 3), starting evening (5 pm), throughout the night and in the morning hours (8 or 9 or 10 am), pitcher temperatures remained below the ambient atmospheric temperatures. The pitcher zones (spots) in six each unopen (sp1-sp3) and open (sp1-sp5) pitchers stayed ≥ 3 degrees below the atmospheric temperatures in the evening-night-morning hours (Figs. 2, 3, S2, S3). This difference (atmospheric temperature-spot) was a maximum of 6.7°C (each) in the unopen pitcher at 4 am (P4U, sp3 - peristome spot) and in open pitcher at 9 pm (P6O, sp5 - lid central spot) (Tables S4, S12). Infrared thermographic profiles demonstrate low temperatures of *Nepenthes* pitcher zones in the evening-night-morning hours (average temperatures, 5 pm to 9 am, unopen pitchers: sp1 20.72, sp2 20.55, sp3 19.95 degrees; open pitchers: sp1 20.96, sp2 20.67, sp3 20.33, sp4 20.24, sp5 19.95 degrees) compared to atmospheric temperatures (average 5 pm to 9 am: 24.13 degrees). In daytime, average atm. temperature was comparatively high, 33.27 degrees (10 am to 4 pm), but pitcher zones were still hotter (average temperatures, 10 am to 4 pm, unopen pitchers: sp1 35.65, sp2 35.50, sp3 36.75 degrees; open pitchers: sp1 34.52, sp2 34.28, sp3 33.47, sp4 33.96, sp5 34.65 degrees). The average humidity in these night and day periods were 77.15% and 50.15%, respectively; on the average 27 units higher in the high evening-night-morning hours (5 pm to 9 am) (Figs. 2, 3; Tables S1-S12). In the high humidity night conditions, moisture from the atmosphere condenses to the entire pitcher surface, particularly on the peristome/lid portions (Fig. 4). In this study, measurements were carried out on non-rainy days; but pitcher peristome may also be wetted by rainfall (Moran and Clarke 2010). Further, the EFN produced by nectaries (Fig. 7) and secretions of glands render wettability to the multiscale peristome cells.

Coincidentally, most *Nepenthes* species are found restricted to perhumid (high humidity index), mountainous regions with warm days and cold and humid nights in their major centres of diversity, *viz.*, Borneo, Sumatra and the Philippines (Moran et al. 2013; Clarke and Moran 2016; Gray et al. 2017). Further Moran and co-workers (2013) demonstrated that prey capture mechanism in carnivorous plants is constrained by climate, *viz.*, *Nepenthes* species employing peristome-based and viscoelastic fluid-based capture are largely restricted to perhumid regions, and species with wax-based mechanism allow successful capture in both perhumid and more seasonal areas. It has also been observed that *Nepenthes* in low light and humidity and high nutrient conditions fails to produce pitchers (Pavlovic et al. 2007). The present data of thermal profiles underline the significance of high humidity in the functioning of *Nepenthes* traps.

In controlled lab conditions (temperature 22°C, humidity 60%) (Fig. 5), sp3 (peristome: average temp. 23.1°C) and sp4 (peristome-lid joint: 22.7°C) showed low temperatures compared to other pitcher zones (sp1 (24.1°C), sp2 (24.2°C), sp5 (24.2°C). These data demonstrate that the peristome surface characteristics favor wettability in *Nepenthes* pitchers (Bohn and Federle 2004; Chen et al. 2016). Further, it is significant to note that even these coldest spots (sp3, sp4) stayed above ambient temperatures (22°C) under these controlled conditions (Fig. 5). This prove the role of high humidity (> 60%, as in night times; Tables S1-S12) and other ecological parameters in maintaining low temperature and wettability of the peristome surface in field conditions. The maximum temperature variation between the spots in controlled condition (22°C/60%) is only 2.1°C, whereas in field these variations are comparatively high (Figs. 2-5). This further evidences the influence of environment on temperature patterns of *Nepenthes* pitchers in the field. Infrared thermographic pictures are showing uniform pattern (blue colour throughout the pitcher) in the nighttime (Figs. 2-4); high humidity in the evening-night-morning conditions favor the adherence moisture to the pitcher surface, lowering its temperature. As the atmospheric temperature rises from morning to noon, moisture evaporates and zones (coloured zones) appear in the pitcher surface (Figs. 2-4).

Thermographic tracking of the pitcher (lid) opening in *N. khasiana* demonstrates the initial ‘wet’ pitcher (coloured blue) in night zone (12.30 am, Fig. 4A), which gradually switches to yellow (relatively hotter) in the daytime (Fig. 4F-I); the peristome surface retains the ‘wetness’ till 10.30 am (Fig. 4H). The pitcher rim (or peristome) in *Nepenthes*, which often extending upwards to the lid forming a distinct neck (Bauer and Federle, 2009), is the hub of prey capture strategies (Figs. 1, 4). The EFN (considered as the major ‘offer’ or ‘reward’; Bauer and Federle 2009; Moran and Clarke 2010) in most *Nepenthes* species is secreted in the extrafloral nectaries located at the inner edge of the peristome, prompting ants and insects to reach towards the cold (wet) region at the deep interior of the pitcher surface (Fig. 7). Bauer and co-workers (2009) illustrated that in wet conditions most ants fell (slipped) into the pitcher when they stepped onto the peristome whereas dry peristome surfaces were not slippery. Simply, peristome wetness and pitcher capture efficiency are in near harmony. Further, experimental peristome wetting in *N. rafflesiana* var. *typica* (Bauer et al. 2008) and *N. bicalcarata* (Bohn and Federle, 2004) demonstrated increased capture efficiency to > 60%. In *N. rafflesiana* var. *typica* pitchers capture efficiency was very high in the evening-night-early morning whereas it was low during the daytime (Bauer et al. 2008). Again, in the evening hours droplets of liquid EFN were noticed on the peristomes of *N. rafflesiana* var. *typica*, and this suggests that EFN secretion plays an important role in peristome wetting (Bauer et al. 2008). Another study showed that the peristome nectaries of *N. rafflesiana* only start secreting after the pitcher has opened, and this coincides with prey capture readiness (Bauer and Federle 2009). Bauer and co-workers further suggested that *Nepenthes* EFN is mainly secreted at times of high humidity when the peristome is already fully wetted, that is, between the evening and early morning (Bauer et al. 2008). Our data of thermal profiles of *Nepenthes* pitchers are further correlating EFN secretion (and peristome wettability) with night time low temperature and high humidity in field conditions.

Chen and co-workers (2016) demonstrated continuous, directional water transport on the surface of *Nepenthes* peristomes. This is due to its multiscale epidermal cells, which optimizes and enhances capillary rise in the transport direction, and prevents backflow by pinning any waterfront that is moving in the reverse direction. This anisotropy (the individual epidermal cells overlap in the same direction, from the outer edge of the peristome inwards) results not only in unidirectional flow (despite the absence of any surface-energy gradient) but also enhances surface wettability (Bauer and Federle 2009; Chen et al. 2016). Similar peristome multiscale structures were also observed in *N. khasiana* (Fig. 7). These unique super-hydrophilic surface properties of the peristome is a critical factor in prey capture. The combination of a hierarchical surface structure and sugary EFN secretions results in an exceptional water-lubricated trapping system. The three dimensional plate-like wax crystals and modified stomata in the hydrophobic slippery zone further prevents the prey from escaping (Hsu et al. 2015; Baby et al. 2017). The hygroscopic EFN secreted at the inner margin of the peristome and hydrophilic surface chemistry associated to surface roughness make the peristome completely covered by a homogenous liquid film which makes it extremely slippery for insects (Bohn and Federle 2004; Bauer et al. 2008; Bauer and Federle 2009; Miguel et al. 2018). In *Nepenthes* pitchers, the waxy surface is almost unwettable, whereas the peristome-lid and digestive surfaces are wettable (Figs. 1, Fig. 7, Baby et al., 2017). The thermographic profiles are displaying the low temperature nighttime zones of peristome-lid portions in *Nepenthes* pitchers and emphasizing their significance in wettability and prey capture. The high wettability of the *Nepenthes* peristome is achieved by a combination of hydrophilicity, surface micro-topography, hygroscopic EFN (Bauer and Federle 2009; Wang et al. 2009) and ecological factors.

### Nepenthes pitchers *versus* thermogenic flowers

Thermoregulation is a trait usually attributed to endothermic animals (mammals, birds). Thermogenesis is also displayed by several plant families; it enhances fragrance production, volatility and has a functional role of providing an energy reward for insect visitors (Seymour et al. 2003; Seymour et al. 2009). Thermogenic flowers are capable of sensing external temperature changes and modulating the rate of heat production controlled by cellular enzymatic processes; and they maintain constant temperature across a wide range of ambient temperatures (Seymour 2001; Seymour et al. 2003). Sacred lotus (*Nelumbo nucifera*), Amazon water lily (*Victoria amazonica*) and *Philodendron selloum* (Araceae) are classic examples of thermogenic plants (Seymour 1997; Lamprecht et al. 1998; Seymour and Matthews 2006). The capacity for heat production varies markedly among thermogenic species, ranging from 2 to 3 degrees to almost 40 degrees above ambient, for a few hours to days and even weeks (for example, inflorescences of *P. selloum*) (Lamprecht et al. 1998; Lamprecht et al. 2002; Grant et al. 2008). Pollinators are able to detect floral thermal patterns, and they prefer this ‘heat reward’ (Seymour 1997; Grant et al. 2008; Harrap et al. 2017). Thermogenesis may also prevent low temperature damage or ensure an optimum temperature for floral development (Grant et al. 2008).

*V. amazonica* is an edible, rhizomatous aquatic plant (Lim et al. 2016) native to SE Colombia, N & E Bolivia and Guyana. Pollination of *V. amazonica* is known to be associated with *Cyclocephala* beetles, and it is known to display floral thermogenicity (Seymour and Matthews 2006). Seymour and Matthews (2006) reported that on the first day the mean floral chamber temperature in *V. amazonica* peaked at 34.7°C at 1940 h and gradually fell during the night to a mean low of 29.3°C at 0620 h; the mean ambient air temperatures decreased from 25.2 to 23.5°C. The grand mean floral chamber temperature was 31.7°C between sunset at 1844 h and sunrise at 0641 h (mean air temperature 24.1°C). On the second night, mean floral temperature was lower at the same times of day, decreasing from 30.2°C (1940 h) to 28.3°C (0620 h); and the mean air temperature decreased from 25.1 to 23.9°C. During the day, temperatures in unhooded flowers rose in response to rising ambient temperature, but remained considerably below it during the hottest part of the day, due to evaporative cooling (Seymour and Matthews 2006). In our infrared thermographic measurements, the base of the floral lobes of *V. amazonica* (sp2) remained hotter almost throughout the 48 h measurements. The flower base (sp1) and top (sp3) petals mostly remained below ambient temperature (Figs. 6, S5, Tables S13, S14). The floral chamber temperatures were not measured in these thermographic experiments.

*N. nucifera* is native to Ukraine, N Iran, far East Russia to tropical Asia and N, NE Australia; its flower is considered as a classic example of beetle pollination (Bernhardt 2000; Davis et al. 2008; Li and Huang 2009). One wild flower of *N. nucifera* produces around 1 million pollen grains, which is an exceptionally high number for a bee-pollinated species (Li and Huang 2009). *N. nucifera* is known to regulate flower temperatures between 30 and 36°C for up to 4 days in environments between 10 and 45°C (Lamprecht et al. 1998; Seymour 1997; Li et al. 2022). In this study (Figs. 6, S6, Table S15), the central floral portions (sp3) of *N. nucifera* remained hotter almost throughout the day; sp3 remained 7.7°C hotter (compared to ambient atmospheric temp.) at 8 am. The inner (sp2) and outer (sp1) floral lobes remained hotter (compared to ambient atmospheric temp.) only in the daytimes (10 am to 3 pm) and (11 am to 4 pm), respectively (Figs. 6C, S6, Table S15).

Recent studies demonstrated that bees are sensitive to floral humidity (von Arx et al. 2012; von Arx 2013; Harrap et al. 2021; Dahake et al. 2022), and on a much larger scale, variations in environmental humidity influence insect behavioral responses (Chown et al. 2011; Enjin 2017). Rands and Whitney (2008) stated that temperature receptors in bee’s antennae are sensitive to temperature variations, and can sense differences as low as 0.25°C. Further, bee responses demonstrated that floral temperature patterns can signal reward location, and therefore they function as floral guides (Harrap et al. 2017; Harrap et al. 2020). In thermogenic plants, floral zones retain temperatures higher than the ambient, thus provides a ‘thermal reward’ and help the pollinator to ‘fly away’. Floral thermogenesis is controlled by enzymatic processes. In contrast, the trapping zones in *Nepenthes* pitchers achieve low temperatures (maximum 6.7°C difference between ambient temperature and *N. khasiana* pitcher spot in the night-morning hours) by physical (peristome microstructures), chemical (EFN) and ecological (humidity, rain) factors. Pollination and prey capture are contrasting ‘life’ and ‘death’ scenarios in nature. The interactions of insects and other arthropods to these pitcher surfaces are to be further investigated considering their sensitivities towards surface (peristome) temperature, humidity, CO2 emission and nectar composition.

Thermogenic plants rely on alternative oxidase (AOX), which is an enzyme in the mitochondria organelle and is a part of the electron transport chain. The reduction of mitochondrial redox potential by AOX increases unproductive respiration. This metabolic process creates an excess of heat which warms thermogenic tissue or organs (Vanlerberghe 2013). Pavlovič and Kocáb (2022) reported high amount of AOX in *Nepenthes* pitcher traps, and discussed its roles in prey attraction, homeostasis of reactive oxygen species, balance between photosynthetic light-dark reactions, digestive physiology and nutrient assimilation. The role of alternative oxidases in pitcher plants are to be further investigated in the context of prey capture.

## Conclusions

*Nepenthes* pitcher peristome and peristome-lid interface acquire low temperatures due to their surface microstructures, secreted nectar and ecological features such as humidity and rain fall. These low temperatures are achieved in evening-night-morning hours when the humidity is high. *Nepenthes* pitcher on opening exposes the (i) peristome surface with microstructures, (ii) peristome fluorescence, (iii) nectaries at the inner side, (iv) peristome and peristome-lid interface with lowest temperatures, (v) releases CO2 to the atmosphere, (vi) toxic metabolites secreted through the nectar and (vi) antifungal metabolites into the pitcher fluid. The peristome epidermal microstructures make it a hydrophilic wettable surface. Wet peristome surface at high humidity (low temperatures) at the night and/or on rainfall mimics ‘aquaplaning’, prevents insect foothold to its surface and makes it extremely slippery.

This study demonstrates that thermal patterns in *Nepenthes* pitchers is supporting their strategy of prey capture. The prey capturing zones (peristome, lid and their intersection) are colder compared to other pitcher regions and ambient temperatures. In contrast, in thermogenic flowers, floral portions are hotter, providing the thermal requirements for the pollinator to fly away and transfer the pollen. Prey capture and pollination are two contrasting natural phenomena, and this study evidences opposite thermal patterns in pitcher and floral organs supporting these events and thereby survival of these plants. Both these phenomena acquired similar attributes (nectar, visual cues, CO2, volatiles) towards their contrasting goals. *Nepenthes* also displays floral traits (pollen, nectar, volatiles) in addition to prey trapping (pitcher) features. Low temperature at the peristome and lid portions of *Nepenthes* pitchers is contradicting the thermal reward offered by thermogenic flowers, but the capture (rates) by these traps are significant favoring the influence of the visual, gaseous and other cues in prey attraction.

## Acknowledgements

We are grateful to Dr. R. Raj Vikraman, Dr. A.A. Prasannakumari and Dr. K.J. Lathan Kumar, Garden Management Division, Jawaharlal Nehru Tropical Botanic Garden and Research Institute, Palode, Thiruvananthapuram for providing us the plant specimens and facilities for the field studies. The field support provided by Mr. Athul Hari S., NFOBC-JRF, Phytochemistry and Phytopharmacology Division during the experiments is also acknowledged.

## Author contributions

SB conceptualized and designed the research, acquired funding; GBS, AJJ performed experiments; SB, GBS, AJJ analyzed the data and prepared the figures; AAZ provided field support; SB wrote the manuscript with contributions from all the authors; All authors read and approved the manuscript.

