## Supplementary Information for "*Nepenthes* pitchers *versus* thermogenic flowers: Thermal patterns and their role in prey capture and pollination"

**Abstract**

Prey capture in *Nepenthes* and pollination in angiosperms are two antithetical events; one designed to trap insects and other arthropods (carnivory) and the other to transfer pollen through pollinators (reproduction). In this study infrared thermography is extensively used to obtain thermal profiles of *Nepenthes* pitchers and thermogenic flowers in field conditions. *N. khasiana* pitchers displayed below ambient temperatures during evening-night-morning hours (5 pm to 8-9-10 am); pitcher spots recorded lowest and highest temperatures as 14.4 (6-7 am) and 45.6$^{o}$C (2 pm), respectively. In the evening-night-morning hours top pitcher spots displayed significant decrease (upto 6.7$^{o}$C) from the ambient temperature. The average humidities in the night and day periods were 77.15% and 50.15%, respectively. Thermographic tracking of the pitcher (lid) opening in *N. khasiana* demonstrated initial 'wet' pitcher in night, which gradually switched to a relatively dry surface in the morning-day times; but the peristome region retained the 'wetness' till 10.30 a.m. The prey capturing zones in *Nepenthes* pitchers (peristome, lid and their intersection) are colder favoring prey capture, whereas in thermogenic flowers, floral portions are hotter, providing the thermal requirements for the pollinator. In thermogenic plants, floral zones assist pollination by offering a 'thermal reward' through enzymatic processes; but *Nepenthes* traps achieve lower temperature spots by physical (surface microstructures), chemical (extrafloral nectar) and ecological (rain, humidity) factors.

*Keywords: Nepenthes*, thermal patterns, prey capture, humidity, thermogenic flowers, pollination.

**Table S1.** Thermographic data of *N. khasiana* unopen pitcher P1U.

| Sl. No. | Pitcher number | Time (h) | Humidity (%) | Temperature (ºC) | sp1 | sp2 | sp3 |
|---|---|---|---|---|---|---|---|
| 1 | P1U | 1800 | 71.0 | 27.8 | 26.9 | 26.6 | 26.6 |
| 2 | P1U | 1900 | 77.0 | 26.1 | 23.8 | 23.3 | 23.0 |
| 3 | P1U | 2000 | 78.1 | 24.9 | 22.2 | 22.1 | 21.7 |
| 4 | P1U | 2100 | 77.3 | 25.2 | 21.5 | 21.4 | 21.0 |
| 5 | P1U | 2200 | 81.0 | 24.7 | 20.7 | 20.5 | 20.3 |
| 6 | P1U | 2300 | 81.0 | 23.4 | 20.3 | 20.2 | 19.8 |
| 7 | P1U | 2400 | 81.0 | 23.0 | 19.6 | 19.5 | 19.2 |
| 8 | P1U | 0100 | 81.0 | 22.8 | 19.1 | 19.1 | 18.7 |
| 9 | P1U | 0200 | 81.0 | 22.6 | 18.5 | 18.6 | 18.2 |
| 10 | P1U | 0300 | 82.0 | 21.7 | 18.1 | 18.0 | 17.6 |
| 11 | P1U | 0400 | 82.0 | 21.7 | 17.6 | 17.6 | 17.3 |
| 12 | P1U | 0500 | 83.5 | 20.6 | 17.8 | 17.7 | 17.4 |
| 13 | P1U | 0600 | 82.0 | 20.4 | 17.7 | 17.8 | 17.5 |
| 14 | P1U | 0700 | 83.0 | 20.9 | 17.6 | 17.8 | 17.5 |
| 15 | P1U | 0800 | 82.0 | 21.2 | 18.4 | 18.9 | 18.6 |
| 16 | P1U | 0900 | 79.0 | 26.3 | 22.3 | 23.4 | 23.3 |
| 17 | P1U | 1000 | 65.5 | 29.8 | 28.8 | 27.3 | 27.2 |
| 18 | P1U | 1100 | 60.6 | 30.7 | 35.3 | 31.2 | 31.1 |
| 19 | P1U | 1200 | 53.0 | 32.9 | 32.7 | 32.4 | 32.3 |
| 20 | P1U | 1300 | 42.0 | 34.5 | 34.1 | 34.0 | 34.0 |
| 21 | P1U | 1400 | 45.0 | 34.8 | 34.3 | 34.0 | 33.9 |
| 22 | P1U | 1500 | 41.5 | 35.4 | 34.6 | 34.8 | 34.0 |
| 23 | P1U | 1600 | 61.5 | 31.4 | 31.4 | 30.9 | 31.0 |
| 24 | P1U | 1700 | 67.5 | 29.7 | 29.6 | 28.8 | 28.7 |

**Table S2.** Thermographic data of *N. khasiana* unopen pitcher P2U.

| Sl. No. | Pitcher number | Time (h) | Humidity (%) | Temperature ($^oC$) | sp1 | sp2 | sp3 |
|---|---|---|---|---|---|---|---|
| 1 | P2U | 1800 | 71.0 | 27.8 | 26.3 | 25.2 | 25.4 |
| 2 | P2U | 1900 | 77.0 | 26.1 | 23.3 | 22.1 | 21.9 |
| 3 | P2U | 2000 | 78.1 | 24.9 | 21.0 | 20.4 | 20.7 |
| 4 | P2U | 2100 | 77.3 | 25.2 | 20.2 | 19.8 | 19.9 |
| 5 | P2U | 2200 | 81.0 | 24.7 | 19.7 | 19.0 | 18.9 |
| 6 | P2U | 2300 | 81.0 | 23.4 | 18.4 | 18.6 | 18.7 |
| 7 | P2U | 2400 | 81.0 | 23.0 | 18.3 | 18.2 | 18.1 |
| 8 | P2U | 0100 | 81.0 | 22.8 | 17.9 | 17.7 | 17.7 |
| 9 | P2U | 0200 | 81.0 | 22.6 | 17.5 | 17.1 | 17.2 |
| 10 | P2U | 0300 | 82.0 | 21.7 | 17.2 | 16.8 | 16.7 |
| 11 | P2U | 0400 | 82.0 | 21.7 | 16.7 | 16.3 | 16.3 |
| 12 | P2U | 0500 | 83.5 | 20.6 | 16.4 | 15.9 | 16.0 |
| 13 | P2U | 0600 | 82.0 | 20.4 | 16.2 | 15.7 | 15.7 |
| 14 | P2U | 0700 | 83.0 | 20.9 | 17.0 | 17.0 | 17.0 |
| 15 | P2U | 0800 | 82.0 | 21.2 | 18.3 | 18.5 | 18.6 |
| 16 | P2U | 0900 | 79.0 | 26.3 | 24.4 | 30.9 | 29.5 |
| 17 | P2U | 1000 | 65.5 | 29.8 | 35.9 | 38.8 | 40.4 |
| 18 | P2U | 1100 | 60.6 | 30.7 | 38.2 | 41.2 | 43.3 |
| 19 | P2U | 1200 | 53.0 | 32.9 | 40.0 | 40.6 | 42.3 |
| 20 | P2U | 1300 | 42.0 | 34.5 | 41.3 | 41.3 | 38.1 |
| 21 | P2U | 1400 | 45.0 | 34.8 | 38.5 | 35.2 | 36.2 |
| 22 | P2U | 1500 | 41.5 | 35.4 | 35.3 | 33.1 | 33.9 |
| 23 | P2U | 1600 | 61.5 | 31.4 | 31.4 | 30.3 | 31.3 |
| 24 | P2U | 1700 | 67.5 | 29.7 | 29.6 | 28.7 | 29.2 |

**Table S3.** Thermographic data of *N. khasiana* unopen pitcher P3U.

| Sl. No. | Pitcher number | Time (h) | Humidity (%) | Temperature (°C) | sp1 | sp2 | sp3 |
|---|---|---|---|---|---|---|---|
| 1 | P3U | 1800 | 71.0 | 27.8 | 27.2 | 26.6 | 25.9 |
| 2 | P3U | 1900 | 77.0 | 26.1 | 24.4 | 23.9 | 23.1 |
| 3 | P3U | 2000 | 78.1 | 24.9 | 22.6 | 22.3 | 21.6 |
| 4 | P3U | 2100 | 77.3 | 25.2 | 21.6 | 21.4 | 20.5 |
| 5 | P3U | 2200 | 81.0 | 24.7 | 21.0 | 20.6 | 19.9 |
| 6 | P3U | 2300 | 81.0 | 23.4 | 20.5 | 20.1 | 19.6 |
| 7 | P3U | 2400 | 81.0 | 23.0 | 19.8 | 19.5 | 19.1 |
| 8 | P3U | 0100 | 81.0 | 22.8 | 19.3 | 19.0 | 18.5 |
| 9 | P3U | 0200 | 81.0 | 22.6 | 18.8 | 18.5 | 17.9 |
| 10 | P3U | 0300 | 82.0 | 21.7 | 18.2 | 18.0 | 17.7 |
| 11 | P3U | 0400 | 82.0 | 21.7 | 18.0 | 17.8 | 17.4 |
| 12 | P3U | 0500 | 83.5 | 20.6 | 17.2 | 17.1 | 16.7 |
| 13 | P3U | 0600 | 82.0 | 20.4 | 16.9 | 16.7 | 16.2 |
| 14 | P3U | 0700 | 83.0 | 20.9 | 17.6 | 17.6 | 17.3 |
| 15 | P3U | 0800 | 82.0 | 21.2 | 18.8 | 19.0 | 18.7 |
| 16 | P3U | 0900 | 79.0 | 26.3 | 26.9 | 29.5 | 29.1 |
| 17 | P3U | 1000 | 65.5 | 29.8 | 31.5 | 33.4 | 36.4 |
| 18 | P3U | 1100 | 60.6 | 30.7 | 33.4 | 33.7 | 36.4 |
| 19 | P3U | 1200 | 53.0 | 32.9 | 34.8 | 36.1 | 36.7 |
| 20 | P3U | 1300 | 42.0 | 34.5 | 37.7 | 39.0 | 43.4 |
| 21 | P3U | 1400 | 45.0 | 34.8 | 39.4 | 39.4 | 43.2 |
| 22 | P3U | 1500 | 41.5 | 35.4 | 40.6 | 42.5 | 44.3 |
| 23 | P3U | 1600 | 61.5 | 31.4 | 33.7 | 34.1 | 33.3 |
| 24 | P3U | 1700 | 67.5 | 29.7 | 30.2 | 30.1 | 29.6 |

**Table S4.** Thermographic data of *N. khasiana* unopen pitcher P4U.

| Sl. No. | Pitcher number | Time (h) | Humidity (%) | Temperature (°C) | sp1 | sp2 | sp3 |
|---|---|---|---|---|---|---|---|
| 1 | P4U | 1800 | 60.0 | 29.0 | 28.5 | 27.3 | 26.2 |
| 2 | P4U | 1900 | 63.0 | 28.2 | 25.4 | 24.5 | 23.6 |
| 3 | P4U | 2000 | 68.0 | 28.0 | 24.6 | 24.4 | 23.4 |
| 4 | P4U | 2100 | 70.0 | 27.5 | 22.8 | 22.3 | 21.1 |
| 5 | P4U | 2200 | 78.0 | 26.0 | 21.2 | 20.8 | 20.0 |
| 6 | P4U | 2300 | 80.0 | 25.2 | 21.1 | 20.6 | 19.8 |
| 7 | P4U | 2400 | 80.0 | 25.0 | 20.0 | 19.4 | 18.5 |
| 8 | P4U | 0100 | 80.0 | 23.5 | 18.9 | 18.6 | 17.9 |
| 9 | P4U | 0200 | 81.0 | 22.0 | 17.8 | 17.1 | 16.5 |
| 10 | P4U | 0300 | 80.0 | 21.5 | 16.7 | 16.3 | 15.7 |
| 11 | P4U | 0400 | 81.0 | 21.3 | 15.7 | 15.3 | 14.6 |
| 12 | P4U | 0500 | 80.0 | 21.2 | 15.6 | 15.4 | 14.9 |
| 13 | P4U | 0600 | 82.0 | 19.0 | 15.4 | 15.2 | 14.7 |
| 14 | P4U | 0700 | 79.0 | 21.0 | 15.2 | 14.9 | 14.4 |
| 15 | P4U | 0800 | 79.0 | 22.0 | 18.3 | 18.8 | 18.3 |
| 16 | P4U | 0900 | 73.0 | 26.0 | 23.4 | 23.3 | 22.1 |
| 17 | P4U | 1000 | 55.0 | 30.7 | 30.0 | 30.2 | 30.0 |
| 18 | P4U | 1100 | 50.0 | 32.5 | 33.6 | 34.2 | 35.1 |
| 19 | P4U | 1200 | 47.0 | 34.5 | 35.9 | 36.7 | 41.0 |
| 20 | P4U | 1300 | 46.0 | 35.0 | 37.4 | 35.1 | 34.1 |
| 21 | P4U | 1400 | 44.0 | 35.5 | 41.5 | 42.9 | 45.4 |
| 22 | P4U | 1500 | 44.0 | 34.2 | 40.1 | 40.6 | 42.8 |
| 23 | P4U | 1600 | 46.9 | 33.8 | 35.3 | 33.9 | 34.1 |
| 24 | P4U | 1700 | 59.4 | 30.7 | 29.9 | 29.0 | 28.6 |

**Table S5.** Thermographic data of *N. khasiana* unopen pitcher P5U.

| Sl. No. | Pitcher number | Time (h) | Humidity (%) | Temperature (°C) | sp1 | sp2 | sp3 |
|---|---|---|---|---|---|---|---|
| 1 | P5U | 1800 | 60.0 | 29.0 | 27.6 | 27.5 | 26.1 |
| 2 | P5U | 1900 | 63.0 | 28.2 | 25.2 | 25.1 | 23.3 |
| 3 | P5U | 2000 | 68.0 | 28.0 | 24.8 | 25.0 | 23.2 |
| 4 | P5U | 2100 | 70.0 | 27.5 | 22.9 | 22.8 | 21.0 |
| 5 | P5U | 2200 | 78.0 | 26.0 | 21.7 | 21.5 | 20.4 |
| 6 | P5U | 2300 | 80.0 | 25.2 | 21.2 | 21.1 | 19.6 |
| 7 | P5U | 2400 | 80.0 | 25.0 | 20.2 | 20.0 | 18.6 |
| 8 | P5U | 0100 | 80.0 | 23.5 | 19.4 | 19.2 | 17.9 |
| 9 | P5U | 0200 | 81.0 | 22.0 | 18.5 | 18.1 | 17.0 |
| 10 | P5U | 0300 | 80.0 | 21.5 | 17.4 | 17.3 | 16.2 |
| 11 | P5U | 0400 | 81.0 | 21.3 | 16.8 | 16.7 | 15.4 |
| 12 | P5U | 0500 | 80.0 | 21.2 | 16.4 | 16.1 | 15.0 |
| 13 | P5U | 0600 | 82.0 | 19.0 | 15.6 | 15.5 | 14.4 |
| 14 | P5U | 0700 | 79.0 | 21.0 | 15.9 | 15.7 | 14.9 |
| 15 | P5U | 0800 | 79.0 | 22.0 | 18.8 | 19.2 | 17.9 |
| 16 | P5U | 0900 | 73.0 | 26.0 | 24.0 | 25.0 | 21.7 |
| 17 | P5U | 1000 | 55.0 | 30.7 | 30.7 | 30.7 | 29.6 |
| 18 | P5U | 1100 | 50.0 | 32.5 | 34.1 | 34.7 | 39.9 |
| 19 | P5U | 1200 | 47.0 | 34.5 | 34.8 | 34.8 | 39.7 |
| 20 | P5U | 1300 | 46.0 | 35.0 | 37.2 | 37.8 | 43.1 |
| 21 | P5U | 1400 | 44.0 | 35.5 | 40.7 | 41.6 | 45.6 |
| 22 | P5U | 1500 | 44.0 | 34.2 | 40.5 | 41.1 | 42.7 |
| 23 | P5U | 1600 | 46.9 | 33.8 | 33.3 | 32.8 | 32.0 |
| 24 | P5U | 1700 | 59.4 | 30.7 | 30.0 | 29.6 | 28.6 |

**Table S6.** Thermographic data of *N. khasiana* unopen pitcher P6U.

| Sl. No. | Pitcher number | Time (h) | Humidity (%) | Temperature (°C) | sp1 | sp2 | sp3 |
|---|---|---|---|---|---|---|---|
| 1 | P6U | 1800 | 60.0 | 29.0 | 28.1 | 27.5 | 26.7 |
| 2 | P6U | 1900 | 63.0 | 28.2 | 25.4 | 24.9 | 24.3 |
| 3 | P6U | 2000 | 68.0 | 28.0 | 24.6 | 24.6 | 23.7 |
| 4 | P6U | 2100 | 70.0 | 27.5 | 23.1 | 22.4 | 21.6 |
| 5 | P6U | 2200 | 78.0 | 26.0 | 22.0 | 21.4 | 20.4 |
| 6 | P6U | 2300 | 80.0 | 25.2 | 20.9 | 20.2 | 19.4 |
| 7 | P6U | 2400 | 80.0 | 25.0 | 20.1 | 19.2 | 18.7 |
| 8 | P6U | 0100 | 80.0 | 23.5 | 19.3 | 18.7 | 18.1 |
| 9 | P6U | 0200 | 81.0 | 22.0 | 18.0 | 17.4 | 16.3 |
| 10 | P6U | 0300 | 80.0 | 21.5 | 17.1 | 16.5 | 16.2 |
| 11 | P6U | 0400 | 81.0 | 21.3 | 16.2 | 15.8 | 15.8 |
| 12 | P6U | 0500 | 80.0 | 21.2 | 16.1 | 15.7 | 15.5 |
| 13 | P6U | 0600 | 82.0 | 19.0 | 15.6 | 15.0 | 14.8 |
| 14 | P6U | 0700 | 79.0 | 21.0 | 15.8 | 15.4 | 15.2 |
| 15 | P6U | 0800 | 79.0 | 22.0 | 19.5 | 20.0 | 19.5 |
| 16 | P6U | 0900 | 73.0 | 26.0 | 25.7 | 26.7 | 25.9 |
| 17 | P6U | 1000 | 55.0 | 30.7 | 30.8 | 36.4 | 41.3 |
| 18 | P6U | 1100 | 50.0 | 32.5 | 37.3 | 33.1 | 31.6 |
| 19 | P6U | 1200 | 47.0 | 34.5 | 34.8 | 33.6 | 33.5 |
| 20 | P6U | 1300 | 46.0 | 35.0 | 38.5 | 35.4 | 36.7 |
| 21 | P6U | 1400 | 44.0 | 35.5 | 37.2 | 35.1 | 35.3 |
| 22 | P6U | 1500 | 44.0 | 34.2 | 36.9 | 35.4 | 35.7 |
| 23 | P6U | 1600 | 46.9 | 33.8 | 33.7 | 31.4 | 31.6 |
| 24 | P6U | 1700 | 59.4 | 30.7 | 30.2 | 28.9 | 29.1 |

**Table S7.** Thermographic data of *N. khasiana* open pitcher P1O.

| Sl. No. | Pitcher number | Time (h) | Humidity (%) | Temperature (°C) | sp1 | sp2 | sp3 | sp4 | sp5 |
|---|---|---|---|---|---|---|---|---|---|
| 1 | P1O | 1800 | 71.0 | 27.8 | 27.2 | 26.7 | 25.6 | 25.7 | 24.7 |
| 2 | P1O | 1900 | 77.0 | 26.1 | 23.7 | 23.4 | 22.8 | 22.5 | 22.4 |
| 3 | P1O | 2000 | 78.1 | 24.9 | 22.5 | 22.2 | 21.8 | 21.7 | 21.7 |
| 4 | P1O | 2100 | 77.3 | 25.2 | 21.6 | 21.4 | 21.1 | 21.0 | 20.9 |
| 5 | P1O | 2200 | 81.0 | 24.7 | 20.7 | 20.6 | 20.4 | 20.2 | 20.2 |
| 6 | P1O | 2300 | 81.0 | 23.4 | 20.0 | 19.7 | 19.5 | 19.5 | 19.3 |
| 7 | P1O | 2400 | 81.0 | 23.0 | 19.7 | 19.6 | 19.5 | 19.2 | 19.2 |
| 8 | P1O | 0100 | 81.0 | 22.8 | 19.1 | 19.0 | 18.8 | 18.6 | 18.5 |
| 9 | P1O | 0200 | 81.0 | 22.6 | 18.5 | 18.3 | 18.1 | 18.0 | 17.8 |
| 10 | P1O | 0300 | 82.0 | 21.7 | 18.3 | 18.0 | 17.8 | 17.7 | 17.4 |
| 11 | P1O | 0400 | 82.0 | 21.7 | 17.9 | 17.7 | 17.5 | 17.3 | 17.2 |
| 12 | P1O | 0500 | 83.5 | 20.6 | 17.5 | 17.3 | 17.0 | 16.9 | 16.9 |
| 13 | P1O | 0600 | 82.0 | 20.4 | 17.3 | 17.1 | 16.9 | 16.9 | 16.7 |
| 14 | P1O | 0700 | 83.0 | 20.9 | 17.9 | 17.9 | 17.7 | 17.5 | 17.4 |
| 15 | P1O | 0800 | 82.0 | 21.2 | 19.0 | 19.2 | 18.8 | 18.7 | 18.5 |
| 16 | P1O | 0900 | 79.0 | 26.3 | 24.3 | 29.5 | 26.1 | 23.2 | 23.7 |
| 17 | P1O | 1000 | 65.5 | 29.8 | 30.1 | 35.0 | 34.4 | 30.2 | 33.9 |
| 18 | P1O | 1100 | 60.6 | 30.7 | 33.2 | 37.2 | 36.6 | 33.7 | 36.2 |
| 19 | P1O | 1200 | 53.0 | 32.9 | 35.3 | 35.1 | 34.7 | 33.1 | 36.1 |
| 20 | P1O | 1300 | 42.0 | 34.5 | 36.4 | 37.0 | 36.1 | 36.6 | 38.2 |
| 21 | P1O | 1400 | 45.0 | 34.8 | 36.7 | 36.5 | 33.1 | 34.0 | 32.9 |
| 22 | P1O | 1500 | 41.5 | 35.4 | 35.7 | 33.4 | 31.7 | 32.8 | 32.5 |
| 23 | P1O | 1600 | 61.5 | 31.4 | 31.5 | 30.6 | 29.5 | 30.0 | 29.9 |
| 24 | P1O | 1700 | 67.5 | 29.7 | 29.1 | 28.5 | 27.5 | 27.7 | 27.0 |

**Table S8.** Thermographic data of *N. khasiana* open pitcher P2O.

| Sl. No. | Pitcher number | Time (h) | Humidity (%) | Temperature (°C) | sp1 | sp2 | sp3 | sp4 | sp5 |
|---|---|---|---|---|---|---|---|---|---|
| 1 | P2O | 1800 | 71.0 | 27.8 | 27.3 | 26.8 | 26.4 | 25.8 | 25.7 |
| 2 | P2O | 1900 | 77.0 | 26.1 | 24.1 | 23.7 | 23.2 | 22.7 | 21.7 |
| 3 | P2O | 2000 | 78.1 | 24.9 | 22.7 | 22.5 | 22.1 | 21.5 | 20.5 |
| 4 | P2O | 2100 | 77.3 | 25.2 | 21.4 | 21.3 | 20.9 | 20.6 | 19.8 |
| 5 | P2O | 2200 | 81.0 | 24.7 | 20.9 | 20.8 | 20.5 | 20.1 | 19.5 |
| 6 | P2O | 2300 | 81.0 | 23.4 | 20.5 | 20.7 | 20.4 | 20.1 | 19.7 |
| 7 | P2O | 2400 | 81.0 | 23.0 | 19.7 | 19.6 | 19.2 | 18.8 | 18.5 |
| 8 | P2O | 0100 | 81.0 | 22.8 | 19.1 | 19.0 | 18.7 | 18.3 | 18.0 |
| 9 | P2O | 0200 | 81.0 | 22.6 | 18.5 | 18.5 | 18.1 | 17.7 | 17.5 |
| 10 | P2O | 0300 | 82.0 | 21.7 | 18.3 | 18.2 | 17.8 | 17.8 | 17.1 |
| 11 | P2O | 0400 | 82.0 | 21.7 | 17.9 | 18.0 | 17.7 | 17.6 | 17.2 |
| 12 | P2O | 0500 | 83.5 | 20.6 | 17.3 | 17.2 | 16.8 | 16.7 | 16.1 |
| 13 | P2O | 0600 | 82.0 | 20.4 | 16.9 | 16.9 | 17.1 | 16.3 | 16.1 |
| 14 | P2O | 0700 | 83.0 | 20.9 | 17.6 | 17.8 | 17.6 | 17.4 | 16.9 |
| 15 | P2O | 0800 | 82.0 | 21.2 | 18.6 | 19.1 | 18.8 | 18.5 | 18.0 |
| 16 | P2O | 0900 | 79.0 | 26.3 | 23.6 | 24.4 | 24.1 | 23.3 | 22.6 |
| 17 | P2O | 1000 | 65.5 | 29.8 | 30.2 | 29.7 | 29.5 | 29.9 | 28.8 |
| 18 | P2O | 1100 | 60.6 | 30.7 | 32.0 | 32.0 | 32.0 | 32.6 | 30.0 |
| 19 | P2O | 1200 | 53.0 | 32.9 | 33.6 | 33.7 | 34.9 | 36.9 | 37.1 |
| 20 | P2O | 1300 | 42.0 | 34.5 | 36.3 | 35.4 | 36.1 | 39.5 | 39.2 |
| 21 | P2O | 1400 | 45.0 | 34.8 | 38.5 | 36.2 | 36.5 | 39.3 | 40.3 |
| 22 | P2O | 1500 | 41.5 | 35.4 | 36.7 | 37.9 | 35.6 | 34.8 | 35.6 |
| 23 | P2O | 1600 | 61.5 | 31.4 | 32.8 | 32.6 | 32.7 | 34.4 | 31.8 |
| 24 | P2O | 1700 | 67.5 | 29.7 | 29.6 | 29.1 | 29.2 | 29.1 | 28.2 |

**Table S9.** Thermographic data of *N. khasiana* open pitcher P3O.

| Sl. No. | Pitcher number | Time (h) | Humidity (%) | Temperature (ºC) | sp1 | sp2 | sp3 | sp4 | sp5 |
|---|---|---|---|---|---|---|---|---|---|
| 1 | P3O | 1800 | 71.0 | 27.8 | 26.6 | 26.2 | 25.7 | 26.0 | 26.0 |
| 2 | P3O | 1900 | 77.0 | 26.1 | 23.3 | 23.1 | 22.7 | 22.9 | 22.8 |
| 3 | P3O | 2000 | 78.1 | 24.9 | 22.1 | 22.0 | 21.8 | 22.0 | 21.9 |
| 4 | P3O | 2100 | 77.3 | 25.2 | 20.9 | 20.8 | 20.7 | 20.9 | 20.8 |
| 5 | P3O | 2200 | 81.0 | 24.7 | 20.8 | 20.7 | 20.6 | 20.8 | 20.8 |
| 6 | P3O | 2300 | 81.0 | 23.4 | 20.1 | 20.0 | 19.8 | 20.0 | 19.9 |
| 7 | P3O | 2400 | 81.0 | 23.0 | 19.7 | 19.6 | 19.6 | 19.7 | 19.7 |
| 8 | P3O | 0100 | 81.0 | 22.8 | 19.1 | 19.0 | 19.0 | 19.0 | 19.0 |
| 9 | P3O | 0200 | 81.0 | 22.6 | 18.4 | 18.4 | 18.3 | 18.3 | 18.3 |
| 10 | P3O | 0300 | 82.0 | 21.7 | 18.1 | 17.9 | 17.9 | 17.9 | 18.0 |
| 11 | P3O | 0400 | 82.0 | 21.7 | 17.7 | 17.7 | 17.6 | 17.7 | 17.8 |
| 12 | P3O | 0500 | 83.5 | 20.6 | 17.1 | 16.9 | 16.9 | 17.0 | 16.9 |
| 13 | P3O | 0600 | 82.0 | 20.4 | 16.7 | 16.6 | 16.5 | 16.6 | 16.6 |
| 14 | P3O | 0700 | 83.0 | 20.9 | 17.4 | 17.5 | 17.4 | 17.5 | 17.5 |
| 15 | P3O | 0800 | 82.0 | 21.2 | 18.5 | 18.6 | 18.5 | 18.6 | 18.7 |
| 16 | P3O | 0900 | 79.0 | 26.3 | 23.0 | 23.0 | 22.7 | 23.0 | 23.1 |
| 17 | P3O | 1000 | 65.5 | 29.8 | 28.2 | 28.2 | 26.3 | 27.9 | 27.1 |
| 18 | P3O | 1100 | 60.6 | 30.7 | 30.9 | 30.7 | 29.6 | 30.9 | 32.2 |
| 19 | P3O | 1200 | 53.0 | 32.9 | 31.6 | 31.9 | 30.8 | 31.3 | 32.8 |
| 20 | P3O | 1300 | 42.0 | 34.5 | 33.2 | 32.9 | 31.9 | 32.3 | 32.9 |
| 21 | P3O | 1400 | 45.0 | 34.8 | 33.7 | 33.0 | 33.5 | 33.0 | 34.2 |
| 22 | P3O | 1500 | 41.5 | 35.4 | 33.5 | 33.2 | 32.6 | 32.4 | 33.1 |
| 23 | P3O | 1600 | 61.5 | 31.4 | 30.4 | 30.4 | 29.4 | 29.7 | 30.4 |
| 24 | P3O | 1700 | 67.5 | 29.7 | 29.3 | 29.0 | 28.0 | 28.4 | 28.6 |

**Table S10.** Thermographic data of *N. khasiana* open pitcher P4O.

| Sl. No. | Pitcher number | Time (h) | Humidity (%) | Temperature (ºC) | sp1 | sp2 | sp3 | sp4 | sp5 |
|---|---|---|---|---|---|---|---|---|---|
| 1 | P4O | 1800 | 60.0 | 29.0 | 28.5 | 27.3 | 26.0 | 26.4 | 26.9 |
| 2 | P4O | 1900 | 63.0 | 28.2 | 25.4 | 24.6 | 24.2 | 24.2 | 24.1 |
| 3 | P4O | 2000 | 68.0 | 28.0 | 25.1 | 24.6 | 24.2 | 24.3 | 24.1 |
| 4 | P4O | 2100 | 70.0 | 27.5 | 23.3 | 22.5 | 22.2 | 22.6 | 22.1 |
| 5 | P4O | 2200 | 78.0 | 26.0 | 22.2 | 21.7 | 21.6 | 21.8 | 21.6 |
| 6 | P4O | 2300 | 80.0 | 25.2 | 21.4 | 20.9 | 20.7 | 21.0 | 20.8 |
| 7 | P4O | 2400 | 80.0 | 25.0 | 20.8 | 20.2 | 20.0 | 20.4 | 20.1 |
| 8 | P4O | 0100 | 80.0 | 23.5 | 19.7 | 19.2 | 18.9 | 19.4 | 19.2 |
| 9 | P4O | 0200 | 81.0 | 22.0 | 18.4 | 17.8 | 17.7 | 17.8 | 17.6 |
| 10 | P4O | 0300 | 80.0 | 21.5 | 17.6 | 17.2 | 17.0 | 17.3 | 17.1 |
| 11 | P4O | 0400 | 81.0 | 21.3 | 16.8 | 16.2 | 16.0 | 16.4 | 16.0 |
| 12 | P4O | 0500 | 80.0 | 21.2 | 16.3 | 15.9 | 15.8 | 15.9 | 15.8 |
| 13 | P4O | 0600 | 82.0 | 19.0 | 15.9 | 15.4 | 15.2 | 15.4 | 15.3 |
| 14 | P4O | 0700 | 79.0 | 21.0 | 15.9 | 15.5 | 15.5 | 15.5 | 15.5 |
| 15 | P4O | 0800 | 79.0 | 22.0 | 19.4 | 19.2 | 18.9 | 19.0 | 18.8 |
| 16 | P4O | 0900 | 73.0 | 26.0 | 24.4 | 24.1 | 22.8 | 22.7 | 22.8 |
| 17 | P4O | 1000 | 55.0 | 30.7 | 30.5 | 29.4 | 28.2 | 28.7 | 28.7 |
| 18 | P4O | 1100 | 50.0 | 32.5 | 35.1 | 34.1 | 34.9 | 36.9 | 44.3 |
| 19 | P4O | 1200 | 47.0 | 34.5 | 35.9 | 34.1 | 34.6 | 34.1 | 36.0 |
| 20 | P4O | 1300 | 46.0 | 35.0 | 36.9 | 34.6 | 33.7 | 34.4 | 35.3 |
| 21 | P4O | 1400 | 44.0 | 35.5 | 40.9 | 42.2 | 35.8 | 36.1 | 37.0 |
| 22 | P4O | 1500 | 44.0 | 34.2 | 38.2 | 40.7 | 40.7 | 35.1 | 42.1 |
| 23 | P4O | 1600 | 46.9 | 33.8 | 33.9 | 33.5 | 33.2 | 31.6 | 33.4 |
| 24 | P4O | 1700 | 59.4 | 30.7 | 30.0 | 29.1 | 28.1 | 28.1 | 28.6 |

**Table S11.** Thermographic data of *N. khasiana* open pitcher P5O.

| Sl. No. | Pitcher number | Time (h) | Humidity (%) | Temperature (ºC) | sp1 | sp2 | sp3 | sp4 | sp5 |
|---|---|---|---|---|---|---|---|---|---|
| 1 | P5O | 1800 | 60.0 | 29.0 | 28.3 | 27.0 | 26.6 | 26.6 | 26.2 |
| 2 | P5O | 1900 | 63.0 | 28.2 | 25.4 | 24.8 | 24.8 | 24.4 | 23.5 |
| 3 | P5O | 2000 | 68.0 | 28.0 | 24.8 | 24.5 | 24.0 | 24.2 | 23.9 |
| 4 | P5O | 2100 | 70.0 | 27.5 | 23.0 | 22.2 | 21.9 | 22.1 | 22.1 |
| 5 | P5O | 2200 | 78.0 | 26.0 | 21.9 | 21.5 | 21.1 | 21.3 | 21.3 |
| 6 | P5O | 2300 | 80.0 | 25.2 | 21.1 | 20.6 | 20.5 | 20.5 | 20.6 |
| 7 | P5O | 2400 | 80.0 | 25.0 | 20.2 | 19.4 | 19.4 | 19.5 | 19.4 |
| 8 | P5O | 0100 | 80.0 | 23.5 | 19.4 | 18.7 | 18.7 | 18.7 | 18.7 |
| 9 | P5O | 0200 | 81.0 | 22.0 | 18.4 | 17.6 | 17.6 | 17.8 | 17.7 |
| 10 | P5O | 0300 | 80.0 | 21.5 | 17.0 | 16.5 | 16.6 | 16.5 | 16.7 |
| 11 | P5O | 0400 | 81.0 | 21.3 | 16.3 | 16.0 | 15.7 | 15.9 | 15.9 |
| 12 | P5O | 0500 | 80.0 | 21.2 | 16.0 | 15.4 | 15.4 | 15.4 | 15.5 |
| 13 | P5O | 0600 | 82.0 | 19.0 | 15.5 | 15.0 | 15.0 | 15.0 | 15.0 |
| 14 | P5O | 0700 | 79.0 | 21.0 | 15.9 | 15.5 | 15.5 | 15.5 | 15.5 |
| 15 | P5O | 0800 | 79.0 | 22.0 | 18.8 | 18.8 | 18.6 | 18.6 | 18.5 |
| 16 | P5O | 0900 | 73.0 | 26.0 | 24.1 | 23.0 | 22.2 | 22.3 | 22.1 |
| 17 | P5O | 1000 | 55.0 | 30.7 | 30.1 | 29.6 | 28.0 | 29.1 | 26.7 |
| 18 | P5O | 1100 | 50.0 | 32.5 | 33.1 | 32.7 | 31.8 | 34.2 | 41.9 |
| 19 | P5O | 1200 | 47.0 | 34.5 | 35.0 | 33.8 | 32.5 | 34.0 | 33.4 |
| 20 | P5O | 1300 | 46.0 | 35.0 | 38.3 | 37.4 | 36.1 | 41.5 | 36.1 |
| 21 | P5O | 1400 | 44.0 | 35.5 | 41.3 | 41.2 | 41.1 | 41.4 | 38.4 |
| 22 | P5O | 1500 | 44.0 | 34.2 | 41.1 | 41.9 | 40.9 | 40.3 | 38.4 |
| 23 | P5O | 1600 | 46.9 | 33.8 | 34.2 | 32.4 | 31.2 | 31.4 | 31.1 |
| 24 | P5O | 1700 | 59.4 | 30.7 | 30.0 | 29.0 | 28.5 | 28.6 | 28.6 |

**Table S12.** Thermographic data of *N. khasiana* open pitcher P6O.

| Sl. No. | Pitcher number | Time (h) | Humidity (%) | Temperature (ºC) | sp1 | sp2 | sp3 | sp4 | sp5 |
|---|---|---|---|---|---|---|---|---|---|
| 1 | P6O | 1800 | 60.0 | 29.0 | 28.0 | 27.2 | 26.6 | 26.4 | 25.6 |
| 2 | P6O | 1900 | 63.0 | 28.2 | 25.5 | 25.0 | 24.7 | 24.3 | 23.1 |
| 3 | P6O | 2000 | 68.0 | 28.0 | 25.4 | 25.0 | 24.6 | 24.5 | 23.2 |
| 4 | P6O | 2100 | 70.0 | 27.5 | 23.4 | 22.8 | 22.3 | 21.9 | 20.8 |
| 5 | P6O | 2200 | 78.0 | 26.0 | 22.0 | 21.8 | 21.6 | 21.3 | 20.5 |
| 6 | P6O | 2300 | 80.0 | 25.2 | 21.7 | 21.0 | 20.7 | 20.5 | 19.6 |
| 7 | P6O | 2400 | 80.0 | 25.0 | 20.4 | 19.8 | 19.5 | 19.5 | 18.7 |
| 8 | P6O | 0100 | 80.0 | 23.5 | 19.6 | 19.0 | 18.7 | 18.7 | 17.6 |
| 9 | P6O | 0200 | 81.0 | 22.0 | 18.6 | 17.8 | 17.4 | 17.2 | 16.5 |
| 10 | P6O | 0300 | 80.0 | 21.5 | 17.4 | 16.8 | 16.5 | 16.3 | 15.5 |
| 11 | P6O | 0400 | 81.0 | 21.3 | 16.7 | 16.3 | 16.0 | 16.1 | 14.7 |
| 12 | P6O | 0500 | 80.0 | 21.2 | 16.4 | 16.0 | 15.7 | 15.7 | 15.0 |
| 13 | P6O | 0600 | 82.0 | 19.0 | 16.2 | 15.8 | 15.5 | 15.4 | 14.6 |
| 14 | P6O | 0700 | 79.0 | 21.0 | 16.5 | 16.0 | 15.4 | 15.2 | 14.7 |
| 15 | P6O | 0800 | 79.0 | 22.0 | 19.8 | 19.6 | 19.3 | 18.8 | 17.8 |
| 16 | P6O | 0900 | 73.0 | 26.0 | 25.5 | 25.7 | 25.1 | 22.5 | 21.5 |
| 17 | P6O | 1000 | 55.0 | 30.7 | 30.6 | 30.1 | 30.2 | 29.9 | 28.7 |
| 18 | P6O | 1100 | 50.0 | 32.5 | 33.8 | 31.8 | 32.5 | 32.2 | 31.6 |
| 19 | P6O | 1200 | 47.0 | 34.5 | 35.6 | 33.7 | 34.0 | 36.8 | 35.0 |
| 20 | P6O | 1300 | 46.0 | 35.0 | 36.4 | 35.3 | 34.5 | 37.3 | 39.0 |
| 21 | P6O | 1400 | 44.0 | 35.5 | 38.2 | 38.2 | 36.0 | 38.1 | 42.0 |
| 22 | P6O | 1500 | 44.0 | 34.2 | 37.5 | 39.2 | 37.3 | 37.0 | 40.5 |
| 23 | P6O | 1600 | 46.9 | 33.8 | 32.7 | 31.2 | 31.1 | 31.0 | 30.7 |
| 24 | P6O | 1700 | 59.4 | 30.7 | 30.1 | 29.2 | 28.4 | 28.4 | 28.7 |

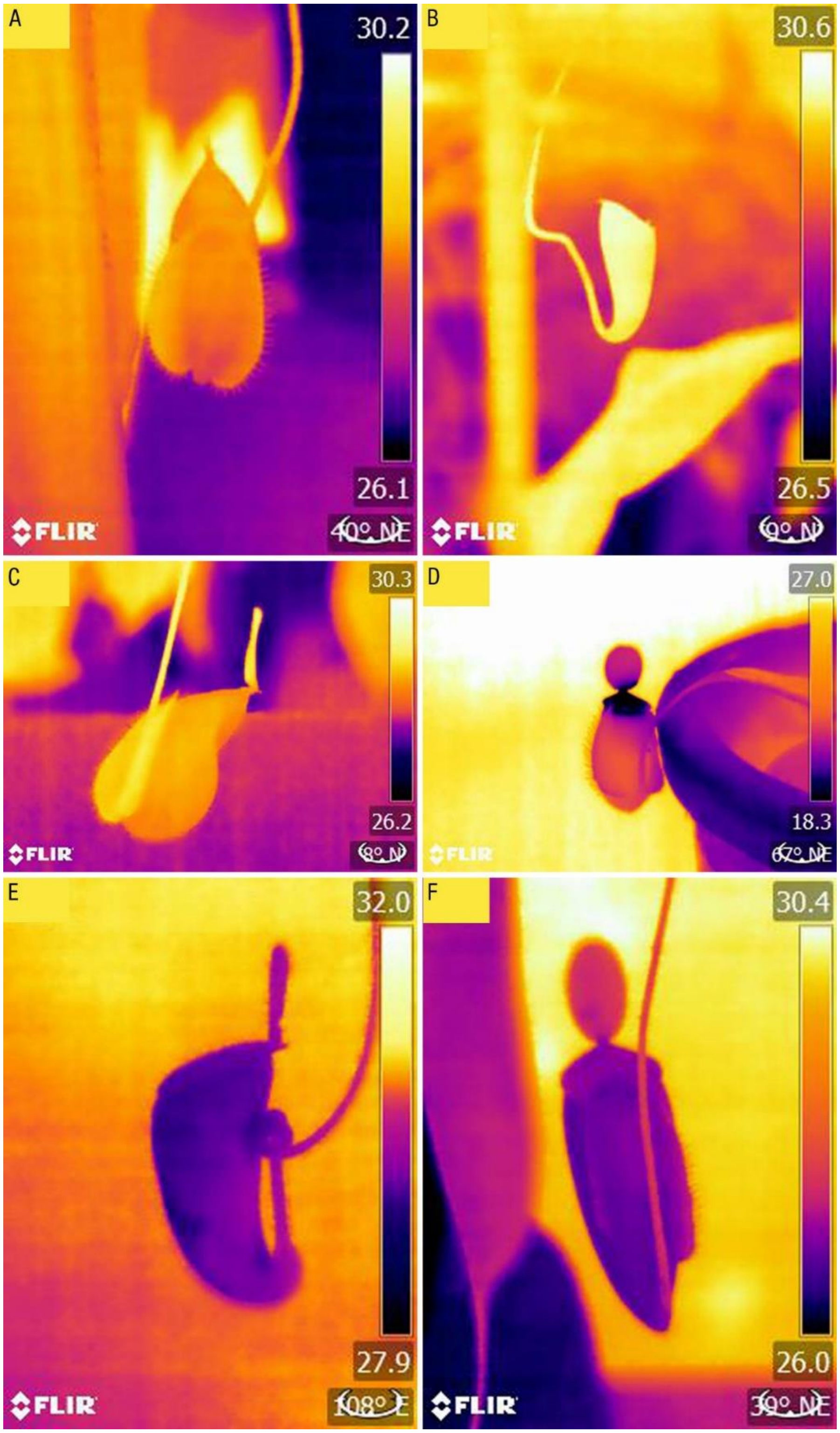


Fig. S1. Thermographic profiles of *Nepenthes* hybrids: (A) hybrid 1 unopen pitcher; (B) hybrid 2 unopen pitcher; (C) hybrid 1 open pitcher; (D) hybrid 2 open pitcher; (E) hybrid 3 open pitcher; (F) hybrid 4 open pitcher.

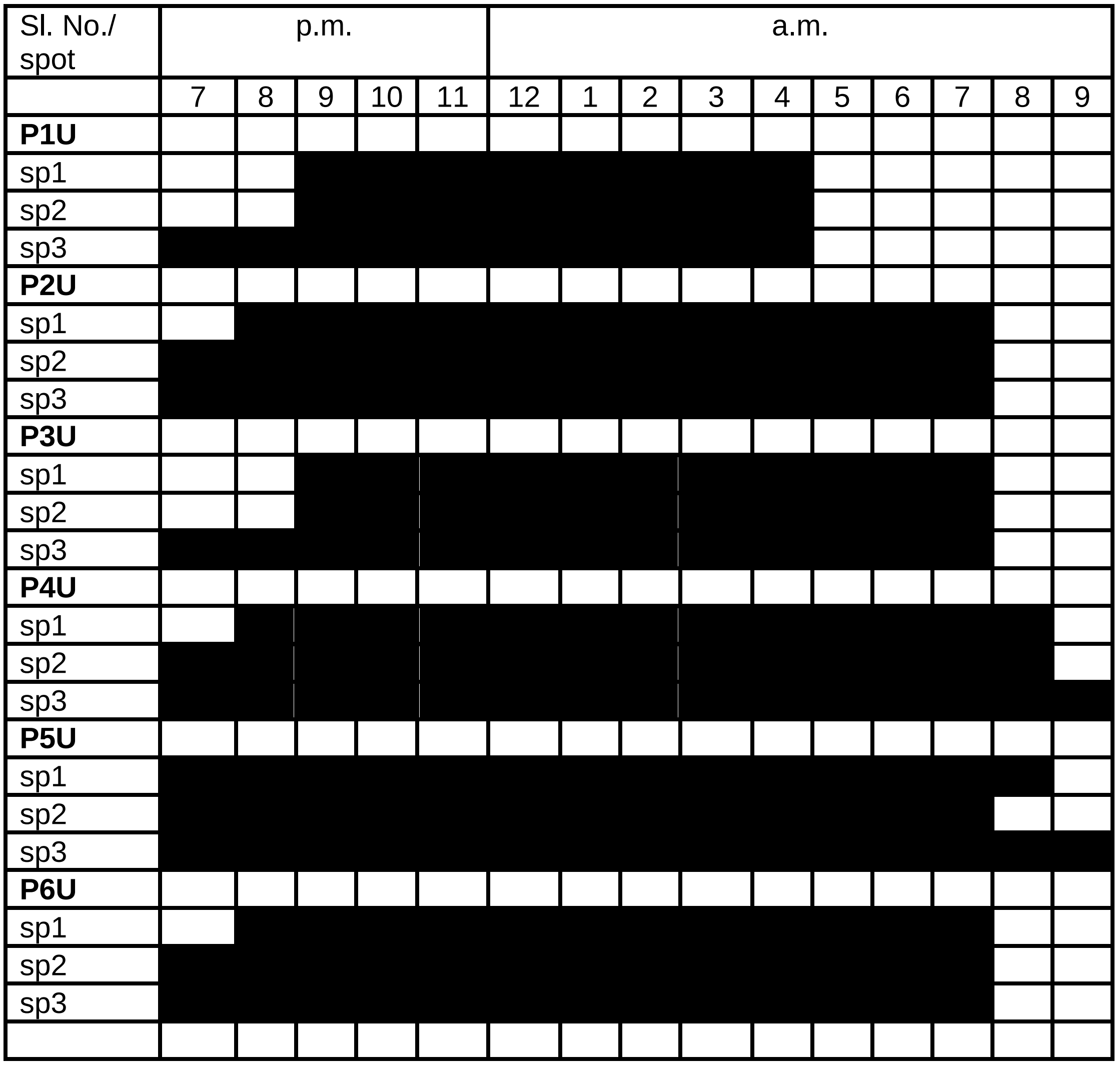


Fig. S2. Nighttime duration (black shaded) in unopen *N. khasiana* pitcher spots (P1U to P6U) at > 3 degrees below the atmospheric temperature.

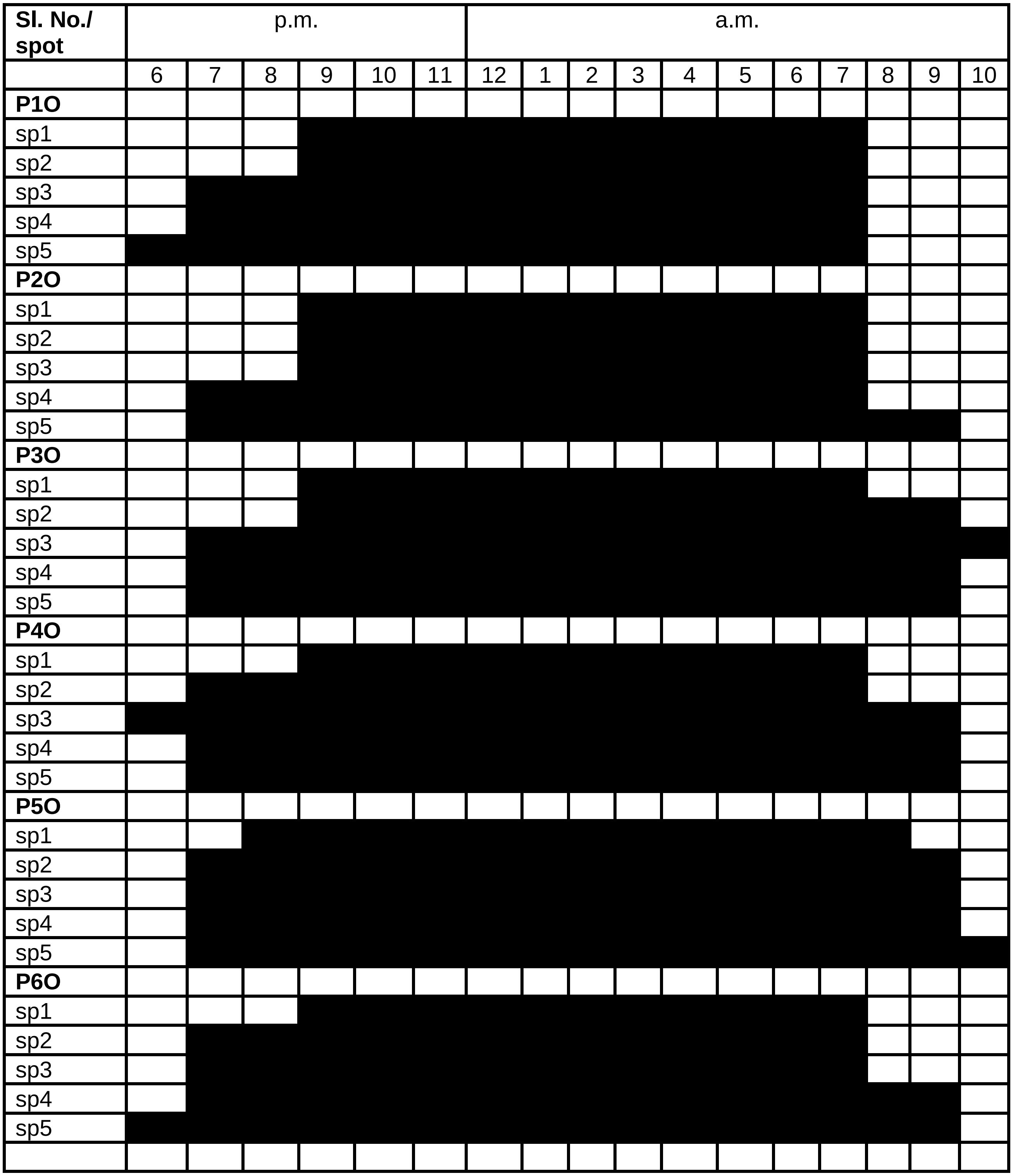


Fig. S3. Nighttime duration (black shaded) in *N. khasiana* open pitcher spots (P1O to P6O) at > 3 degrees below the atmospheric temperature.

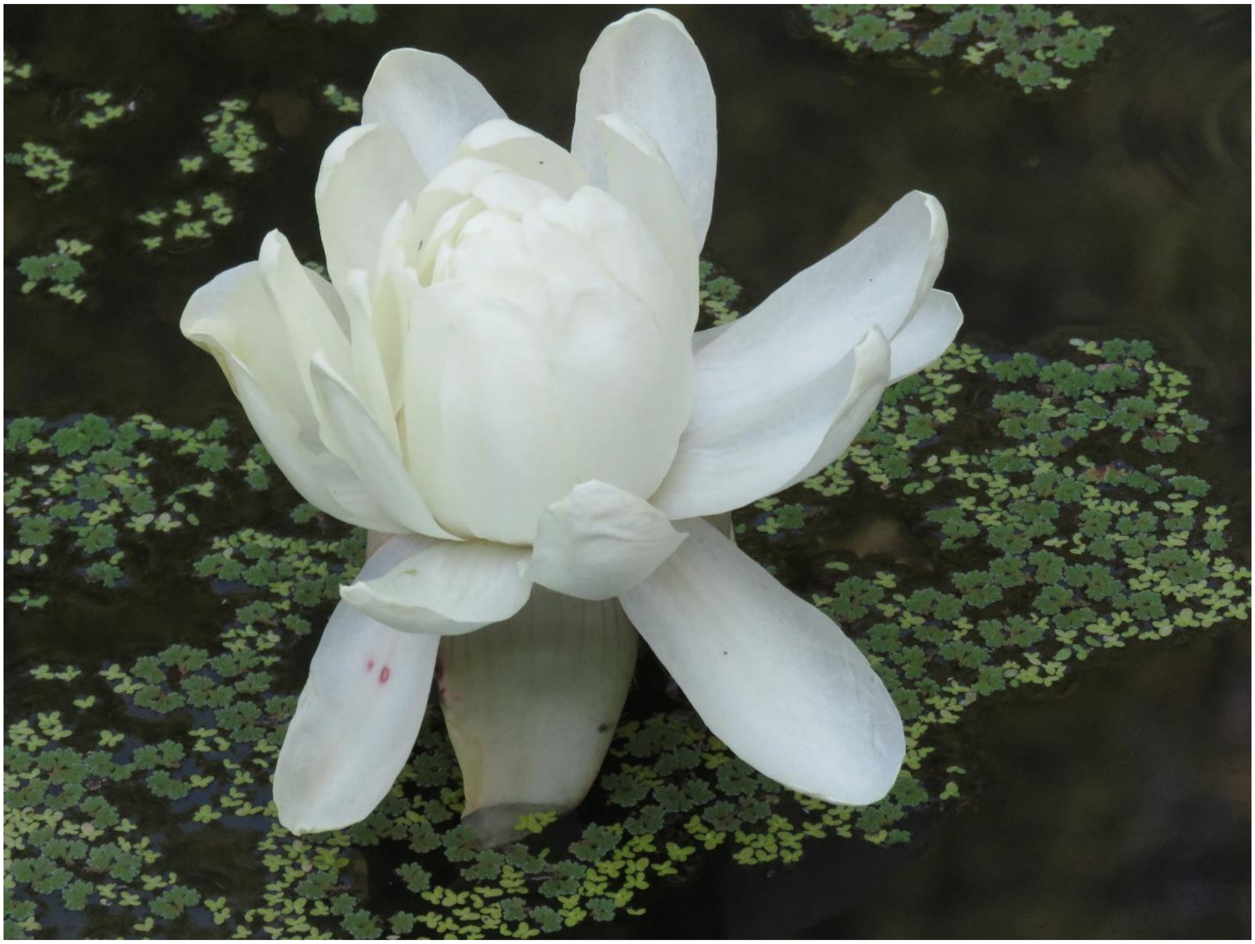

Fig. S4. *Victoria amazonica* flower.

**Table S13.** Thermographic data of *Victoria amazonica* (January 2019).

| Sl. No. | Species | Time | Humidity (%) | Temperature ( $^{o}$C) | sp1 | sp2 | sp3 |
|---|---|---|---|---|---|---|---|
| 1 | *V. amazonica* | 16.00 | 61.5 | 31.4 | - | 28.4 | 28.2 |
| 2 | *V. amazonica* | 17.00 | 67.5 | 29.7 | - | 29.3 | 26.7 |
| 3 | *V. amazonica* | 18.00 | 75.0 | 27.0 | - | 30.1 | 25.1 |
| 4 | *V. amazonica* | 19.00 | 80.0 | 25.8 | 28.6 | 29.6 | 25.2 |
| 5 | *V. amazonica* | 20.00 | 82.0 | 25.7 | 25.3 | 28.0 | 23.9 |
| 6 | *V. amazonica* | 21.00 | 81.0 | 25.2 | 23.4 | 27.9 | 22.9 |
| 7 | *V. amazonica* | 22.00 | 82.4 | 24.9 | 23.1 | 28.0 | 22.3 |
| 8 | *V. amazonica* | 23.00 | 82.2 | 24.3 | 23.2 | 27.0 | 21.8 |
| 9 | *V. amazonica* | 24.00 | 82.8 | 23.7 | 22.0 | 26.6 | 21.3 |
| 10 | *V. amazonica* | 01.00 | 80.8 | 23.8 | 21.7 | 25.5 | 20.8 |
| 11 | *V. amazonica* | 02.00 | 83.3 | 22.6 | 21.3 | 25.2 | 20.3 |
| 12 | *V. amazonica* | 03.00 | 82.5 | 22.6 | 21.0 | 24.6 | 19.6 |
| 13 | *V. amazonica* | 04.00 | 83.0 | 21.9 | 20.7 | 24.2 | 19.1 |
| 14 | *V. amazonica* | 05.00 | 81.6 | 22.0 | 19.6 | 23.6 | 19.1 |
| 15 | *V. amazonica* | 06.00 | 83.5 | 21.2 | 19.9 | 23.8 | 18.4 |
| 16 | *V. amazonica* | 07.00 | 85.3 | 20.5 | 20.1 | 23.6 | 19.4 |
| 17 | *V. amazonica* | 08.00 | 80.0 | 23.8 | 21.6 | 26.4 | 22.7 |
| 18 | *V. amazonica* | 09.00 | 78.0 | 26.8 | 27.1 | 32.1 | 31.3 |
| 19 | *V. amazonica* | 10.00 | 83.0 | 26.0 | 29.5 | 34.0 | 30.1 |

**Table S14.** Thermographic data of *V. amazonica* (January 2022).

| Sl. No. | Species | Time | Humidity (%) | Temperature ( $^{o}$C) | sp1 | sp2 | sp3 |
|---|---|---|---|---|---|---|---|
| 1 | *V. amazonica* | 16.00 | 65.0 | 33.0 | - | 28.6 | 29.5 |
| 2 | *V. amazonica* | 17.00 | 85.0 | 28.8 | - | 28.6 | 26.0 |
| 3 | *V. amazonica* | 18.00 | 84.0 | 27.2 | - | 28.9 | 25.4 |
| 4 | *V. amazonica* | 19.00 | 84.0 | 27.0 | 25.0 | 29.0 | 24.6 |
| 5 | *V. amazonica* | 20.00 | 83.5 | 26.7 | 25.1 | 27.7 | 24.2 |
| 6 | *V. amazonica* | 21.00 | 84.3 | 25.9 | 25.0 | 28.2 | 23.2 |
| 7 | *V. amazonica* | 22.00 | 83.0 | 26.0 | 23.6 | 27.3 | 22.8 |
| 8 | *V. amazonica* | 23.00 | 83.6 | 26.0 | 23.4 | 26.8 | 22.2 |
| 9 | *V. amazonica* | 24.00 | 83.7 | 25.7 | 23.3 | 26.1 | 22.3 |
| 10 | *V. amazonica* | 01.00 | 83.8 | 25.4 | 22.5 | 25.5 | 22.0 |
| 11 | *V. amazonica* | 02.00 | 82.4 | 25.4 | 22.6 | 24.0 | 21.2 |
| 12 | *V. amazonica* | 03.00 | 83.7 | 24.4 | 22.3 | 23.7 | 20.9 |
| 13 | *V. amazonica* | 04.00 | 82.0 | 24.0 | 21.6 | 23.8 | 20.3 |
| 14 | *V. amazonica* | 05.00 | 82.3 | 23.4 | 21.5 | 23.8 | 20.0 |
| 15 | *V. amazonica* | 06.00 | 83.0 | 23.0 | 21.3 | 22.7 | 20.1 |
| 16 | *V. amazonica* | 07.00 | 83.2 | 23.6 | 21.7 | 23.8 | 20.8 |
| 17 | *V. amazonica* | 08.00 | 84.0 | 24.3 | 23.3 | 26.3 | 22.9 |
| 18 | *V. amazonica* | 09.00 | 79.5 | 28.5 | 24.5 | 29.3 | 25.7 |
| 19 | *V. amazonica* | 10.00 | 79.0 | 28.9 | 26.4 | 29.1 | 26.3 |
| 20 | *V. amazonica* | 11.00 | 72.0 | 31.7 | 28.4 | 30.1 | 29.2 |
| 21 | *V. amazonica* | 12.00 | 64.1 | 31.8 | 29.1 | 31.0 | 30.8 |
| 22 | *V. amazonica* | 13.00 | 57.0 | 36.6 | 30.7 | 31.3 | 33.7 |
| 23 | *V. amazonica* | 14.00 | 64.3 | 31.5 | 29.4 | 30.5 | 30.4 |
| 24 | *V. amazonica* | 15.00 | 68.0 | 30.3 | 27.9 | 27.7 | 28.3 |
| 25 | *V. amazonica* | 16.00 | 76.0 | 28.5 | 28.1 | 28.8 | 27.8 |
| 26 | *V. amazonica* | 17.00 | 74.5 | 30.3 | 27.7 | 28.3 | 26.9 |
| 27 | *V. amazonica* | 18.00 | 76.0 | 28.5 | 26.2 | 27.0 | 24.9 |
| 28 | *V. amazonica* | 19.00 | 81.2 | 27.0 | 25.6 | 28.8 | 24.2 |
| 29 | *V. amazonica* | 20.00 | 80.0 | 27.1 | 25.9 | 31.0 | 23.5 |
| 30 | *V. amazonica* | 21.00 | 83.1 | 26.2 | 25.9 | 30.0 | 25.2 |
| 31 | *V. amazonica* | 22.00 | 81.8 | 26.0 | 25.7 | 28.8 | 23.5 |
| 32 | *V. amazonica* | 23.00 | 82.8 | 26.1 | 25.2 | 27.9 | 23.5 |
| 33 | *V. amazonica* | 24.00 | 81.5 | 26.0 | 24.1 | 26.8 | 22.8 |
| 34 | *V. amazonica* | 01.00 | 84.1 | 25.0 | 24.1 | 26.4 | 23.0 |
| 35 | *V. amazonica* | 02.00 | 83.6 | 24.9 | 24.0 | 26.0 | 22.1 |
| 36 | *V. amazonica* | 03.00 | 83.0 | 26.0 | 24.3 | 25.4 | 22.7 |
| 37 | *V. amazonica* | 04.00 | 83.8 | 24.6 | 23.7 | 25.3 | 22.0 |

| | | | | | | | |
|---|---|---|---|---|---|---|---|
| 38 | *V. amazonica* | 05.00 | 83.0 | 24.3 | 23.6 | 24.7 | 22.7 |
| 39 | *V. amazonica* | 06.00 | 82.5 | 23.3 | 23.5 | 23.5 | 21.4 |
| 40 | *V. amazonica* | 07.00 | 84.9 | 23.7 | 23.6 | 23.8 | 22.6 |
| 41 | *V. amazonica* | 08.00 | 82.7 | 26.2 | 24.9 | 25.7 | 24.5 |
| 42 | *V. amazonica* | 09.00 | 78.0 | 29.1 | 27.8 | 30.8 | 29.3 |
| 43 | *V. amazonica* | 10.00 | 65.5 | 31.6 | 29.8 | 34.5 | 30.1 |
| 44 | *V. amazonica* | 11.00 | 75.5 | 29.4 | 28.3 | 30.7 | 30.1 |
| 45 | *V. amazonica* | 12.00 | 73.0 | 30.6 | 28.5 | 32.2 | 31.8 |
| 46 | *V. amazonica* | 13.00 | 65.5 | 30.6 | 30.3 | 33.7 | 32.7 |
| 47 | *V. amazonica* | 14.00 | 65.5 | 34.2 | 38.0 | 37.1 | 40.2 |
| 48 | *V. amazonica* | 15.00 | 75.7 | 31.5 | 28.2 | 30.9 | 30.5 |

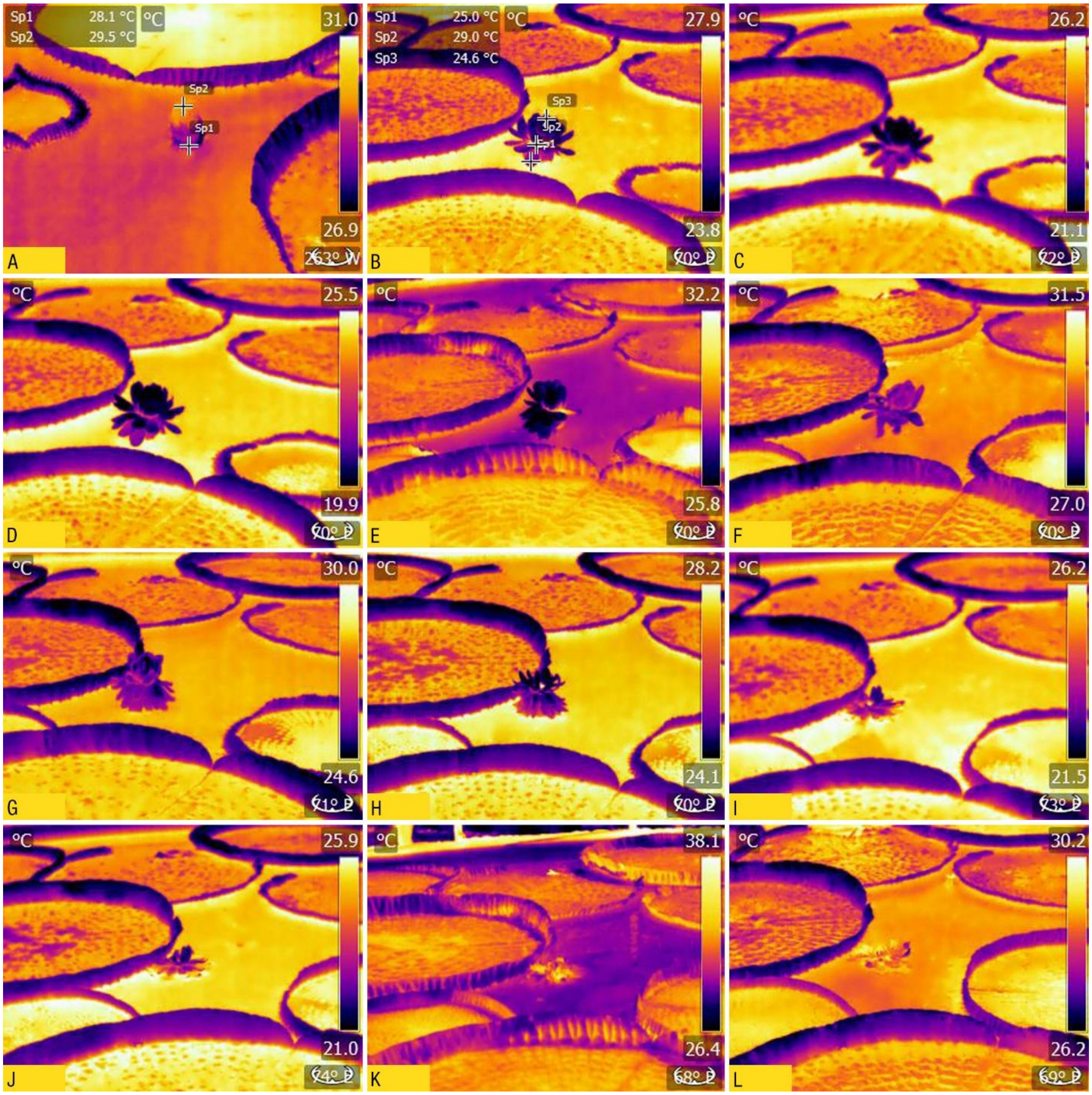


Fig. S5. Thermographic profiles of *V. amazonica* (A) flower bud above the level of water (1600); (B) flower bloomed, three clear spots (sp1, sp2, sp3) visible (1900); (C & D) flower showing three spots but intensity of sp2 decreased (0200 & 0600); (E) flower is slightly pink and started to fold (1000); (F) flower in completely pink, partially opened, sp2 is visible but with low intensity (1600); (G) flower fully opened and sp2 is of low intensity (1800); (H) flower top is more opened and a hot central portion is visible (1900); (I) flower is partially floating in water, central region is visible (0200); (J) flower is partially floating, central region visible with low intensity (0600); (K) flower floats above water not spot differentiation visible (1000); (L) flower almost immersed in water with some portions floating above, nearly dead state (1600).

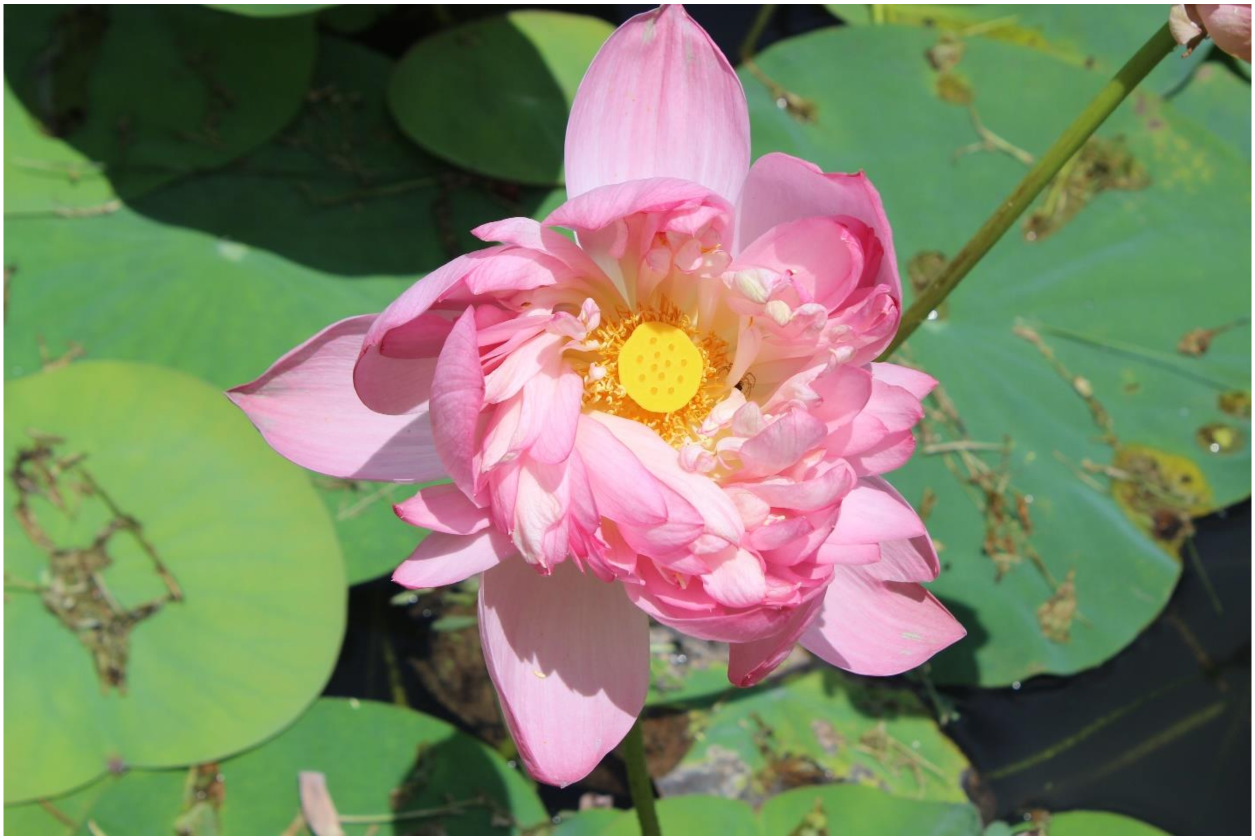

Fig. S6. *Nelumbo nucifera* flower.

**Table S15.** Thermographic data of *Nelumbo nucifera* (March 2021).

| Sl. No. | Species | Time (h) | Humidity (%) | Temperature (°C) | sp1 | sp2 | sp3 |
|---|---|---|---|---|---|---|---|
| 1 | *N. nucifera* | 1800 | 60.0 | 29.0 | 27.3 | 25.2 | 30.3 |
| 2 | *N. nucifera* | 1900 | 63.0 | 28.2 | 24.6 | 26.7 | 29.2 |
| 3 | *N. nucifera* | 2000 | 68.0 | 28.0 | 24.2 | 26.3 | 30.2 |
| 4 | *N. nucifera* | 2100 | 70.0 | 27.5 | 22.8 | 25.7 | 30.6 |
| 5 | *N. nucifera* | 2200 | 78.0 | 26.0 | 22.0 | 24.7 | 29.9 |
| 6 | *N. nucifera* | 2300 | 80.0 | 25.2 | 21.3 | 24.5 | 28.6 |
| 7 | *N. nucifera* | 2400 | 80.0 | 25.0 | 20.5 | 23.0 | 27.3 |
| 8 | *N. nucifera* | 0100 | 80.0 | 23.5 | 19.3 | 21.4 | 25.4 |
| 9 | *N. nucifera* | 0200 | 81.0 | 22.0 | 18.2 | 20.2 | 24.4 |
| 10 | *N. nucifera* | 0300 | 80.0 | 21.5 | 17.1 | 18.3 | 23.0 |
| 11 | *N. nucifera* | 0400 | 81.0 | 21.3 | 16.7 | 18.2 | 21.4 |
| 12 | *N. nucifera* | 0500 | 80.0 | 21.2 | 16.1 | 18.3 | 20.9 |
| 13 | *N. nucifera* | 0600 | 82.0 | 19.0 | 15.7 | 16.6 | 20.4 |
| 14 | *N. nucifera* | 0700 | 79.0 | 21.0 | 16.7 | 17.4 | 18.9 |
| 15 | *N. nucifera* | 0800 | 79.0 | 22.0 | 22.6 | 23.1 | 29.7 |
| 16 | *N. nucifera* | 0900 | 73.0 | 26.0 | 24.5 | 24.8 | 30.1 |
| 17 | *N. nucifera* | 1000 | 55.0 | 30.7 | 27.7 | 28.8 | 33.3 |
| 18 | *N. nucifera* | 1100 | 50.0 | 32.5 | 35.2 | 37.4 | 39.1 |
| 19 | *N. nucifera* | 1200 | 47.0 | 34.5 | 34.9 | 38.3 | 38.6 |
| 20 | *N. nucifera* | 1300 | 46.0 | 35.0 | 36.9 | 42.7 | 39.2 |
| 21 | *N. nucifera* | 1400 | 44.0 | 35.5 | 38.8 | 39.1 | 37.6 |
| 22 | *N. nucifera* | 1500 | 44.0 | 34.2 | 35.9 | 36.4 | 34.8 |
| 23 | *N. nucifera* | 1600 | 46.9 | 33.8 | 31.4 | 32.1 | 32.7 |
| 24 | *N. nucifera* | 1700 | 59.4 | 30.7 | 27.9 | 27.9 | 29.1 |

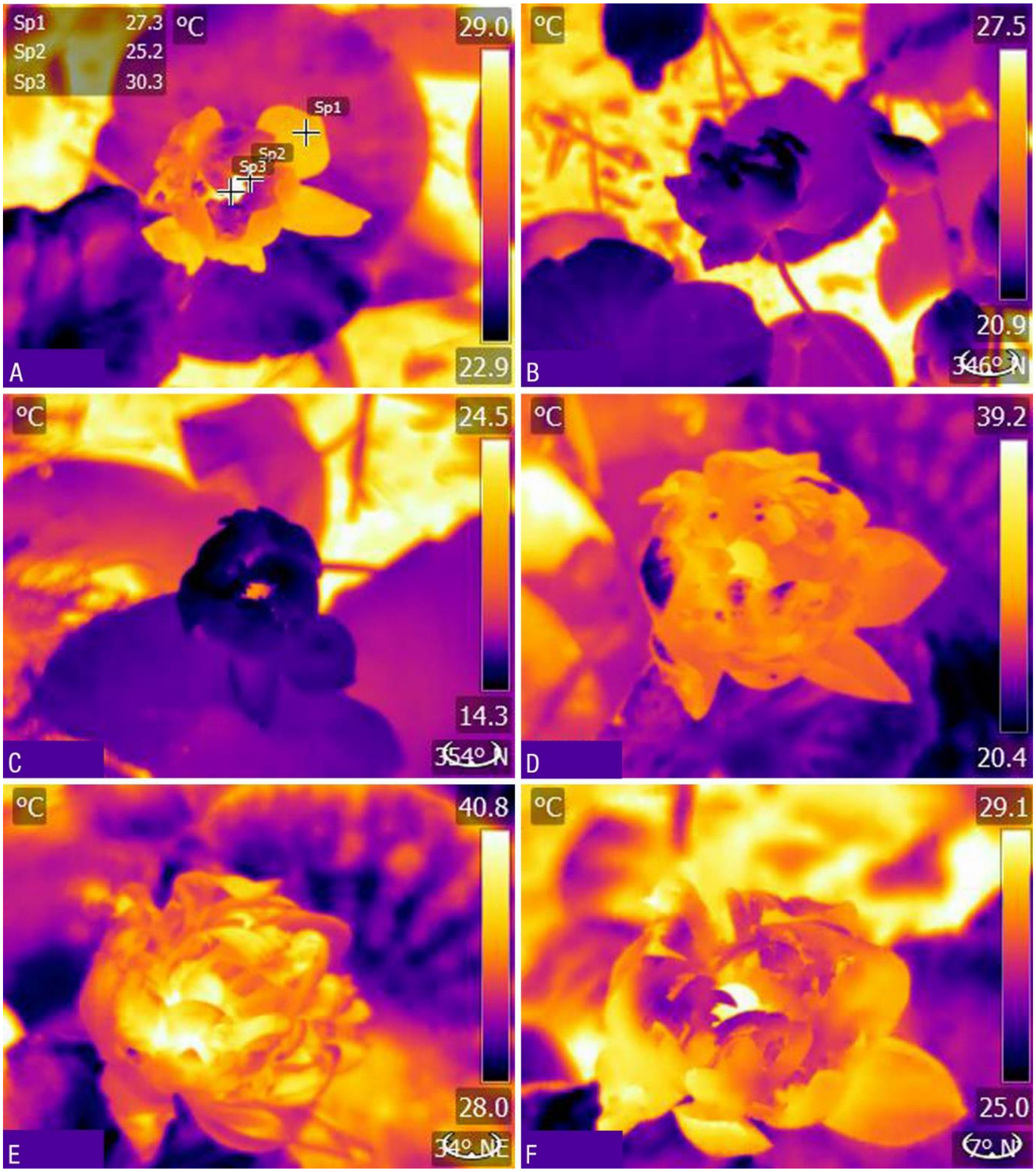


Fig. S7. Thermographic profiles of *N. nucifera* (A) fully bloomed flower in open pond, three regions clearly spotted (1800); (B) flower partially closed, three regions not clearly visible (2200); (C) flower partially opened, regions begins to appear (0600); (D) flower completely opened, all regions clearly spottable (1000); (E) flower, totally opened and is moisture free (1100); (F) flower began to close, all regions visible (1800).
